# The centriculum, a membrane reticulum that surrounds *C. elegans* centrosomes, may serve as a microtubule filter

**DOI:** 10.1101/2025.08.16.670680

**Authors:** Richa Maheshwari, Mohammad M Rahman, Abigail Ruddick, Seth Drey, Robert S Mirabello, Orna Cohen-Fix

**Affiliations:** The Laboratory of Biochemistry and Genetics, National Institute of Diabetes and Digestive and Kidney Disease, National Institutes of Health, Bethesda MD, 20892, USA; University of Oregon, Department of Biology, Eugene, OR 97403, USA; University of Louisville School of Medicine, Louisville KY, 40202, USA

**Keywords:** centriculum, centrosome, *C. elegans*, microtubules, SPD-5

## Abstract

Centrosomes are cytoplasmic microtubule-nucleating structures. They are considered membraneless organelles, but in several cell types they are surrounded by ER-derived membrane. In *C. elegans* early embryos, this membrane forms a dense membrane reticulum, named the centriculum, that affects centrosome structure and microtubule nucleating capacity. The centriculum is adjacent to the centrosome’s pericentriolar material (PCM) and to abundant, short peri-centrosomal microtubules. Here we show that when microtubule abundance is reduced, centriculum and PCM size decrease, while the density of the PCM protein SPD-5 increases, suggesting that the PCM can be compacted. We further show that centriculum size is determined by microtubules, and that the centriculum likely limits the length of peri-centrosomal microtubules. Finally, we find that the centriculum is more porous where spindle microtubules pass compared to where astral microtubules pass. These data are consistent with the centriculum serving as a microtubule filter, blocking the extension of most, but not all, centrosome-nucleated microtubules. Finally, if microtubule-centriculum collisions result in microtubule catastrophe, the filter function of the centriculum could also explain the high concentration of soluble tubulin at the *C. elegans* centrosome.

**Summary Statement:** The centriculum is a centrosome-associated membrane reticulum in *C. elegans* embryos. This study shows that its size depends on microtubules and that it may act as a filter, limiting microtubule growth.

## Introduction

Centrosomes are cytoplasmic structures that nucleate microtubules, polymers made of α- and β-tubulin heterodimers that form linear, side-by-side, protofilaments, creating a hollow microtubule tube (reviewed in (Aljiboury & Hehnly, 2023)). Microtubules are polar; centrosome-nucleated microtubules have their minus (α-tubulin facing) ends embedded in the centrosome, and their plus (β-tubulin facing) ends extending away from the centrosome. Microtubules are dynamic structures that grow and shrink mainly at their plus end in a manner that is dependent on soluble tubulin concentrations and aided by microtubule associated proteins, such as motor proteins, microtubule polymerases and severing proteins, and more (reviewed in (Gudimchuk & McIntosh, 2021). In dividing cells, centrosomes nucleate microtubules that either form the mitotic spindle or extend to the cell cortex in the form of astral microtubules (reviewed in (Hoffmann, 2021)).

At the core of centrosomes are two centrioles; they are surrounded by a pericentriolar material (PCM), a proteinaceous structure whose properties are just beginning to emerge (reviewed in (Magescas *et al*, 2019; Pimenta-Marques & Bettencourt-Dias, 2020; Woodruff *et al*, 2014). The PCM is rich in coiled-coil proteins that form a lattice, or scaffold, onto which additional PCM proteins are recruited. One of the main components of the PCM in *C. elegans* is SPD-5 (Hamill *et al*, 2002), which is functionally homologous to Centrosomin/Cnn in *Drosophila* (Megraw *et al*, 2001; Megraw *et al*, 1999; Timothy *et al*, 1999) and CDK5RAP2 in vertebrates (Fong *et al*, 2008). The PCM also includes γ-tubulin (TBG-1 in *C. elegans*), which, along with associated proteins that form the γ-tubulin ring complex (γ -TuRC), serves to nucleate microtubules ((Moritz *et al*, 1995; Zheng *et al*, 1995) reviewed in (Gao *et al*, 2024)). At the onset of mitosis, the PCM increases in size in a process known as centrosome maturation ((Khodjakov & Rieder, 1999) reviewed in (Vasquez-Limeta & Loncarek, 2021)). The expansion of the *C. elegans* PCM, as measured by SPD-5 accumulation, involves the incorporation of soluble SPD-5 throughout the volume of the SPD-5 lattice (Laos *et al*, 2015; Woodruff *et al*, 2015). This process is facilitated by phosphorylation of SPD-5 by the polo-like kinase PLK-1 (Cabral *et al*, 2019; Nakajo *et al*, 2022; Rios *et al*, 2024; Woodruff *et al*., 2015; Wueseke *et al*, 2016). PLK-1 phosphorylation of SPD-5 is independently required to promote γ-tubulin recruitment to the PCM (Ohta *et al*, 2021). The role of PLK-1 in centrosome maturation is conserved throughout eukaryotes (Conduit *et al*, 2015; Lane & Nigg, 1996; Lee & Rhee, 2011).

Centrosomes are considered membraneless organelles, although there are numerous examples of endoplasmic reticulum (ER)-derived membrane accumulation adjacent to the centrosome in *Drosophila*, sea urchin, the fish medaka, mammalian cultured cells, and *C. elegans* (Araújo *et al*, 2023; Bergman *et al*, 2015; Bobinnec *et al*, 2003; Diaz *et al*, 2019; Harris, 1975; Karabasheva & Smyth, 2019; Rollins & Blankenship, 2023; Schlaitz, 2014; Terasaki, 2000; Waterman-Storer *et al*, 1993). This peri-centrosomal membrane was analyzed in the *C. elegans* 1-cell embryo by volume electron microscopy at 9 nm isotropic resolution and found to be made of a dense membrane reticulum, leading to its name the centriculum (Maheshwari *et al*, 2023). While the structure of peri-centrosomal membranes in other organisms has yet to be examined at this resolution, the presence of the ER curvature-inducing proteins Rtn1 and ReepB at peri-centrosomal membranes of *Drosophila* embryos (Diaz *et al*., 2019) suggests a similar structure. In *C. elegans*, down-regulation of atlastin, which forms ER-ER junctions (Hu & Rapoport, 2016), led to an increase in centriculum size (Maheshwari *et al*., 2023). This was accompanied by an increase in both centrosome size and the amount of PCM material (as measured by the area and total amount of fluorescently tagged PCM proteins), and increased microtubule-nucleating capacity (Maheshwari *et al*., 2023). Likewise, in *Drosophila* embryos, alteration to ER structure, including prominent changes to the membrane surrounding centrosomes, caused defects in centrosome and spindle structure (Araújo *et al*., 2023; Rollins & Blankenship, 2023). Thus, at least in these systems, centrosome function is affected by the membrane adjacent to it, indicating that the centrosome may not be as membraneless as previously assumed.

It is well documented that during *C. elegans* early embryogenesis, the majority of microtubules nucleated from the surface of the centrosome terminate in the vicinity of the centrosome (for example, see (Albertson, 1984; Redemann *et al*, 2017) and Fig. 1A; defined here as peri-centrosomal microtubules). A similar accumulation of short microtubules around centrosomes is seen in embryos of medaka (Kiyomitsu *et al*, 2024), *Drosophila* (Rollins & Blankenship, 2023) and sea urchin (Xie *et al*, 2025), all of which also have centrosome-associated ER membranes. The *C. elegans* centrosome is also able to accumulate soluble tubulin at a concentration that is 10-fold higher than in the cytoplasm (Baumgart *et al*, 2019). The precise mechanism that facilitates this accumulation is not known. A condensate of SPD-5 with additional PCM proteins was shown to concentrate soluble tubulin *in vitro* (Woodruff *et al*, 2017), although not to the extent observed *in vivo* (Baumgart *et al*., 2019). The centriculum provides an attractive mechanism for these two phenomena: First, peri-centrosomal microtubule accumulation could reflect collisions of a subset of microtubules with centriculum membranes that are in their path and impede their elongation. In this way, the centriculum would act as a microtubule filter, limiting the number of microtubules that can fully elongate. This could reduce the competition for cytoplasmic tubulin pools and/or prevent the formation of a poorly functioning spindle. Second, if these collided microtubules terminate at the centriculum and undergo catastrophe (akin to the catastrophe that occurs when microtubules encounter the plasma membrane (Bouvrais *et al*, 2021)), this could provide a source for the exceedingly high soluble tubulin concentration at the centrosome (Baumgart *et al*., 2019). We therefore set out to determine the spatial relationship between the centrosome, centriculum and peri-centrosomal microtubules, and to examine whether the centriculum functions as a microtubule filter.

**Figure 1:**
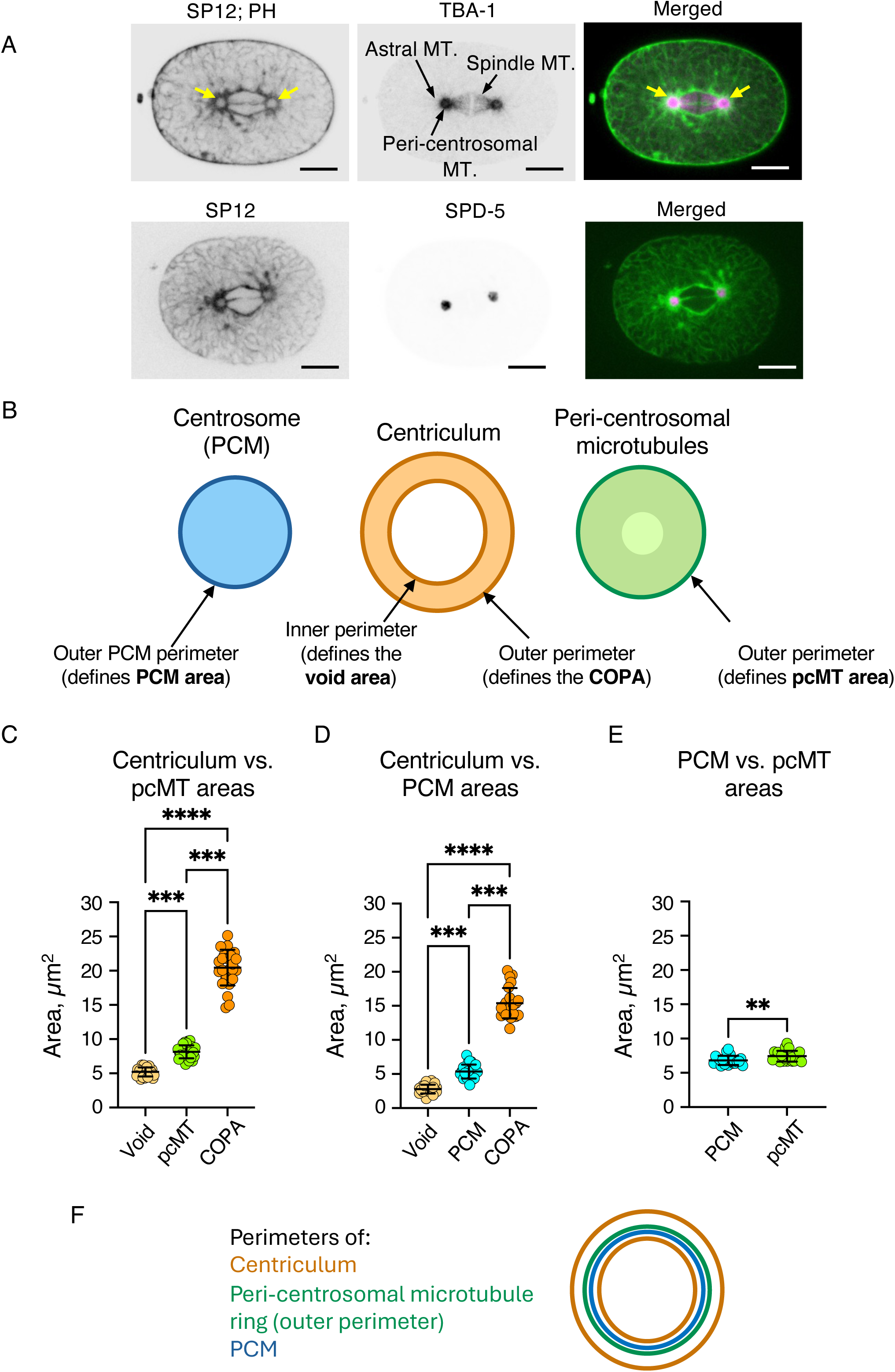
The spatial relationship between the centrosome, centriculum and peri-centrosomal microtubules. **A)** Top panel: a *C. elegans* 1-cell embryo (strain OCF193) at metaphase expressing GFP::SP12 and PH domain::GFP (ER and plasma membrane markers, respectively; left panel and in green in merged panel) and TBA-1::RFP (microtubules; middle panel and in magenta in merged panel). Three types of microtubules are indicated: astral microtubules, which are nucleated at the centrosome and extend toward the cortex, spindle microtubules, which extend toward the chromosomes (not shown, but are located in the gap between the two half spindles), and peri-centrosomal microtubules, which accumulate in the vicinity of the centrosome but do not extend much past it. The two centricula flanking the spindle are indicated by yellow arrows in the left and merged images. Lower panel: *C. elegans* embryo (strain OCF176) at metaphase expressing GFP::SP12 (ER marker; green in merged panel) and GFP::SPD-5 (PCM marker; magenta in merged panel). Scale bar=10 µm. **B)** Schematic representation of the centrosome (PCM), with its area (light blue) defined by its outer perimeter (dark blue line); the centriculum, outlined by its inner and outer perimeters (dark orange lines; the area encompassed by the inner perimeter is defined as the centriculum “void”, and the area enclosed by the outer perimeter of the centriculum is defined as the centriculum outer perimeter area, or COPA); and peri-centrosomal microtubule (pcMT) area (light green), defined by its outer perimeter (dark green). **C**, **D and E)** Quantification of void area (light orange), peri-centrosomal microtubules (green), PCM (blue) and COPA (dark orange) in 1-cell embryos from control RNAi treated worms. The strains and their relevant markers were as follows: Panel C: OCF181 (mCherry::SP12; GFP::TBA-2; n=26); Panel D: OCF176 (mCherry::SP12; GFP::SPD-5; n=22 for void area, n=24 for PCM, and n=23 for COPA; Panel E: OCF247 (GFP::SPD-5; TBA-1:: tagRFP) n=20. Please refer to Fig. S1B for comparison with strain OCF240 (RFP::SPD-5; GFP::TBA-2). Statistical analyses: Panel C: p=0.0001 for void area vs pcMT area; p<0.0001 for void area vs COPA; p=0.0001 for pcMT area vs COPA. Panel D: p=0.0004 for void area vs PCM area; p<0.0001 for void area vs COPA; p=0.0002 for PCM area vs COPA. These analyses were done using Anova Kruskal-Wallis test. Panel E: p=0.0043 using Anova Mann-Whitney test. Error bars indicate mean and standard deviation. **F)** Schematic representation of the spatial relationship between the perimeters of the centriculum outer and inner perimeter (orange), peri-centrosomal microtubules (green) and the PCM (blue).

## Results

### The spatial relationship between the centrosome, centriculum and microtubules

In our previous study ((Maheshwari *et al*., 2023), we showed that the centriculum surrounds the centrosome and the peri-centrosomal microtubules (Fig. 1A), but the exact spatial relationship between the three was not quantified. To do so, we examined these three structures at metaphase of the *C. elegans* 1-cell embryo, when parental genomes are still encased in separate pronuclei (Fig. 1A). In 1-cell embryos, centricula become visible as soon as centrosomes separate (while still associated with the male pronucleus), and both centricula and centrosomes increase in size throughout the cell cycle (Hamill *et al*., 2002; Maheshwari *et al*., 2023). Consequently, size measurements must be done at the same cell cycle stage, which in our case is at metaphase. At this stage, the two pronuclei are adjacent to each other, and the membrane interface between them is still present (Fig. 1A). Microtubules that are nucleated from the outer portion of the PCM (O’Toole *et al*, 2012) extend outwards, towards the cortex (as astral microtubules), or chromosome (as spindle microtubules). However, only a fraction of these microtubules extends well beyond the centrosome (Redemann *et al*., 2017), creating an accumulation of peri-centrosomal microtubules (Fig. 1A, top panel). To determine the spatial relationship between the centrosome, the centriculum and the peri-centrosomal microtubules, we used confocal imaging to capture the areas encompassed by the structures’ perimeter(s) (Fig. 1B), visualized at their largest cross section; as these structures are nested within each other, the plane of their largest cross-section is shared. For the centrosome, we define “PCM area” as the area occupied by SPD-5, the outermost component of the PCM (Magescas *et al*., 2019). For the centriculum, visualized using fluorescently tagged SP12 (Poteryaev *et al*, 2005), we defined two areas: the area encompassed by the inner perimeter of the centriculum, referred to as the centriculum “void area”, and the area encompassed by the outer perimeter of the centriculum, referred to as “centriculum outer perimeter area”, or “COPA” (note that the area of the centriculum itself is the difference between COPA and the centriculum void area). For peri-centrosomal microtubules, visualized using fluorescently tagged α or β tubulin, we defined the area encompassed by the outer perimeter of the fluorescent tubulin signal as “the peri-centrosomal microtubule area” (referred to in figures as “pcMT area”, for simplicity). Although peri-centrosomal microtubules often appear as a ring due to the exclusion of microtubules from the center of the centrosome (Strome *et al*, 2001), the difference in fluorescence between the ring and its “hole” is only about 10-15% (likely due to the high concentration of soluble tubulin), making “hole” size determination somewhat ambiguous. Thus, for our discussion, the area occupied by peri-centrosomal microtubules includes the “hole”. The comparison of areas defined by the perimeters of the three structures was done using pairwise combinations; imaging all three structures at once was not feasible due to a weak signal when proteins were fused to blue/cyan fluorescent proteins. Parenthetically, we observed that the absolute size of various structures sometimes differed between strains, even when they were otherwise isogenic (see, for example, the sizes of the void area and COPA in Figs. 1C and D). Thus, the relative sizes of two structures were determined by comparisons within the same strain. In the next set of experiments, worms were treated with control RNAi so that the measurements could be compared to the various RNAi treatments, described below.

The results of the pairwise combinations are shown in Figs. 1C-E and Fig. S1B. As expected, the area defined by the outer perimeter of the centriculum, COPA, was the largest. The area defined by the inner perimeter of the centriculum, the void area, was the smallest. The area defined by the outer perimeter of peri-centrosomal microtubules was greater than the area of the PCM (using two different fluorescently tagged protein combinations, see Fig. 1E and Fig S1B), and both were greater than the centriculum void area, suggesting that they abut or even extend into the centriculum. For microtubules, we know this to be the case based on EM tomography data (Maheshwari *et al*., 2023). Based on these data, an approximation of the overall spatial relationship between the centrosome, centriculum and peri-centrosomal microtubules in wild type cells is shown in Fig. 1F.

### Decrease in microtubule number or stability results in a smaller centriculum and PCM compression

The proximity between the centrosome, centriculum and peri-centrosomal microtubules raised the question of size dependency: Which of the three structures determines the dimensions of one or both other structures? We previously showed that reducing the levels of KLP-7, a microtubule depolymerase whose down-regulation leads to enhanced microtubule outgrowth from centrosomes (Srayko *et al*, 2005), resulted in a larger centriculum and a larger SPD-5 area (Maheshwari *et al*., 2023). This suggested that microtubule stability or abundance affects centriculum and/or PCM size. To further examine this possibility, we examined the consequences of reduced microtubule elongation or nucleation. To this end, we down-regulated by RNAi either *zyg-9* or *tbg-1*. ZYG-9 is a microtubule polymerase and a homologue of *Xenopus* XMAP215 and human ch-TOG. It localizes to centrosomes and the mitotic spindle (Gard & Kirschner, 1987; Matthews *et al*, 1998). In *Xenopus*, XMAP215 has been shown to bind free tubulin dimers and catalyze their addition to the growing microtubule plus-end (Brouhard *et al*, 2008). TBG-1 is the *C. elegans* γ-tubulin, which localizes to centrosomes and facilitates microtubule nucleation (note that in *C. elegans*, TBG-1/ γ-tubulin is not essential for microtubule nucleation at centrosomes (Hamill *et al*., 2002; Hannak *et al*, 2002; Strome *et al*., 2001). Both conditions are expected to reduce the amount of peri-centrosomal microtubules (as determined by the total fluorescence of tubulin around the centrosome), as is indeed the case (Fig. S1C). The area occupied by peri-centrosomal microtubules was also reduced, as was the size of the centriculum (Figs. 2A-C; *zyg-9* and *tbg-1* RNAi data are compared to control RNAi data also shown in Fig. 1C). These results further support our conclusion that microtubule abundance and/or stability affects centriculum size.

**Figure 2:**
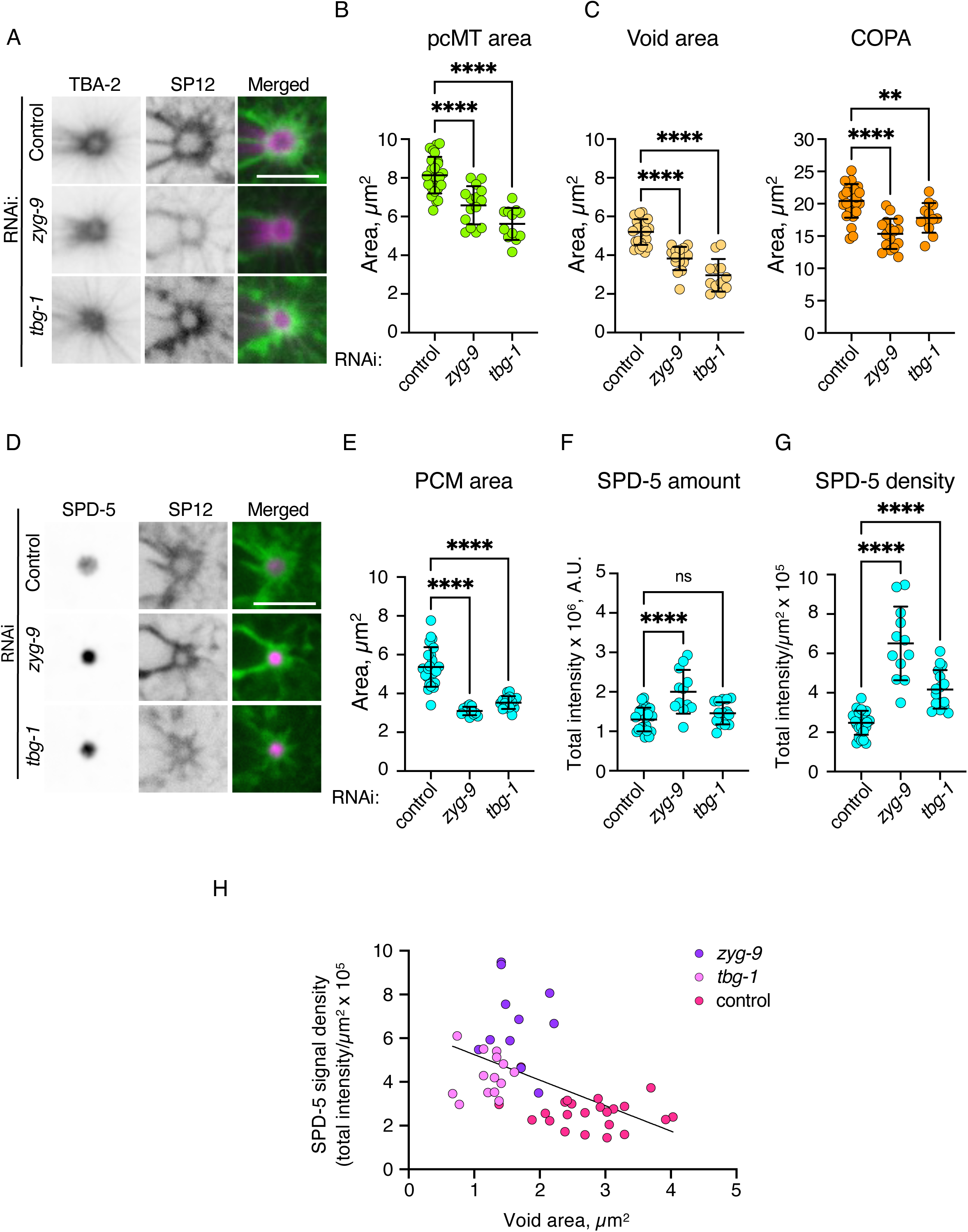
Decrease in microtubule abundance or stability leads to a smaller centriculum and compression of the PCM. **A)** Representative images of centricula from 1-cell embryos (OCF181) at metaphase expressing the SP12 ER marker fused to mCherry (mCherry::SP12; green in merged images) and α-tubulin fused to GFP (GFP::TBA-2; magenta in merged images), from worms treated with RNAi against a control (top row), *zyg-9* (middle row) or *tbg-1* RNAi (bottom row). Scale bar=5 μm. **B and C)** Quantification of GFP::TBA-2 peri-centrosomal microtubule area, void area and COPA, from embryos treated as shown in panel A. Panel B: n=26, 15, and 12 for control, *zyg-9* and *tbg-1* RNAi treatments, respectively. p< 0.0001 for control vs *zyg-9* and control vs *tbg-1* RNAi treatments using ordinary one-way ANOVA. Panel C: n=26, 16 and 12 for control, *zyg-9* and *tbg-1* RNAi treatments, respectively. For void area: p< 0.0001 for control vs *zyg-9* and control vs *tbg-1* RNAi treatments using Kruskal-Wallis test. For COPA: p< 0.0001 for control vs *zyg-9* RNAi and p=0.0072 for control vs *tbg-1* RNAi using ordinary one-way ANOVA. Error bars represent mean and standard deviation. Note that the values for control RNAi are the same as shown in Fig. 1C. **D)** Representative images of centricula and PCM from 1-cell embryos (OCF176) at metaphase expressing mCherry::SP12 (green in merged images) and the SPD-5 PCM protein fused to GFP (GFP::SPD-5; magenta in merged images), from worms treated with RNAi against a control (top row), *zyg-9* (middle row) or *tbg-1* (bottom row). Scale bar=5 μm. **E, F and G)** Quantification of GFP::SPD-5 area, amount and density (determined by SPD-5 total intensity divided (panel F) by SPD-5 area (panel E)) from embryos treated as shown in panel D. n=24, 12, and 16 for control, *zyg-9* and *tbg-1* RNAi treatments, respectively. For panel E: p<0.0001 for control vs *zyg-9* and for control vs *tbg-1* RNAi treatments using ordinary one-way ANOVA. Note that the control RNAi values for SPD-5 area are the same as in Fig. 1D. Panel F: p=<0.0001 for control vs *zyg-9* and p=0.3290 for control vs *tbg-1* RNAi treatments using ordinary one-way ANOVA. Panel G: p=<0.0001 for control vs *zyg-9* and for control vs *tbg-1* RNAi treatments using ordinary one-way ANOVA. Error bars indicate mean and standard deviation. **H)** GFP::SPD-5 density (as in panel G) as a function of void area (Fig. S1D) following control (magenta), *zyg-9* (purple), or *tbg-1* (pink) RNAi treatments. Linear regression was performed through all the datapoints. R^2^=0.2624; slope was significantly different from 0 (p=0.0002).

The reduction in centriculum size following *zyg-9* or *tbg-1* RNAi led us to examine what happens to SPD-5 under these conditions. Using a strain expressing the SP12 ER marker fused to mCherry and GFP::SPD-5, we observed that down-regulation of *zyg-9* or *tbg-1* was accompanied by a decrease in centriculum size, as before (Fig. S1D), and in PCM size (Figs. 2D and E). Interestingly, while the area occupied by SPD-5 decreased, there was no loss of SPD-5 from the PCM (Fig. 2F), and SPD-5’s density in the PCM increased (Fig. 2G). Overall, the increase in SPD-5 density was inversely proportional to size of the centriculum void area (Fig. 2H; R^2^=0.2624 for all data points). This result suggests that centrosome size is affected by either microtubule abundance and/or the size of the centriculum. This further suggests that the PCM is compressible (see Discussion).

### The *spd-5* expansion mutant, in the presence of wild type *spd-5*, reduces overall PCM size

The possibility that the centriculum affects centrosome size was intriguing, because the centrosome is considered a membraneless organelle. In this scenario, centriculum size is determined by the abundance and/or stability of microtubules, and the PCM expands to the boundaries of the centriculum (Fig. 3Ai). Alternatively, however, centriculum size could be determined by PCM size, while PCM size is determined by microtubules abundance and/or stability (Fig. 3Aii). This latter scenario seemed less likely, because PCM maturation is thought to be microtubule-independent (Hannak *et al*., 2002; Khodjakov & Rieder, 1999). Nonetheless, we needed to establish whether, in our system, centriculum size was determined by the PCM or microtubules. To do so, we needed a condition that differentially alters the areas occupied by the PCM vs peri-centrosomal microtubules. We could then determine with which of the two centriculum size correlates. This condition was met by using a *spd-5* allele that is defective in expanding the SPD-5 lattice.

**Figure 3:**
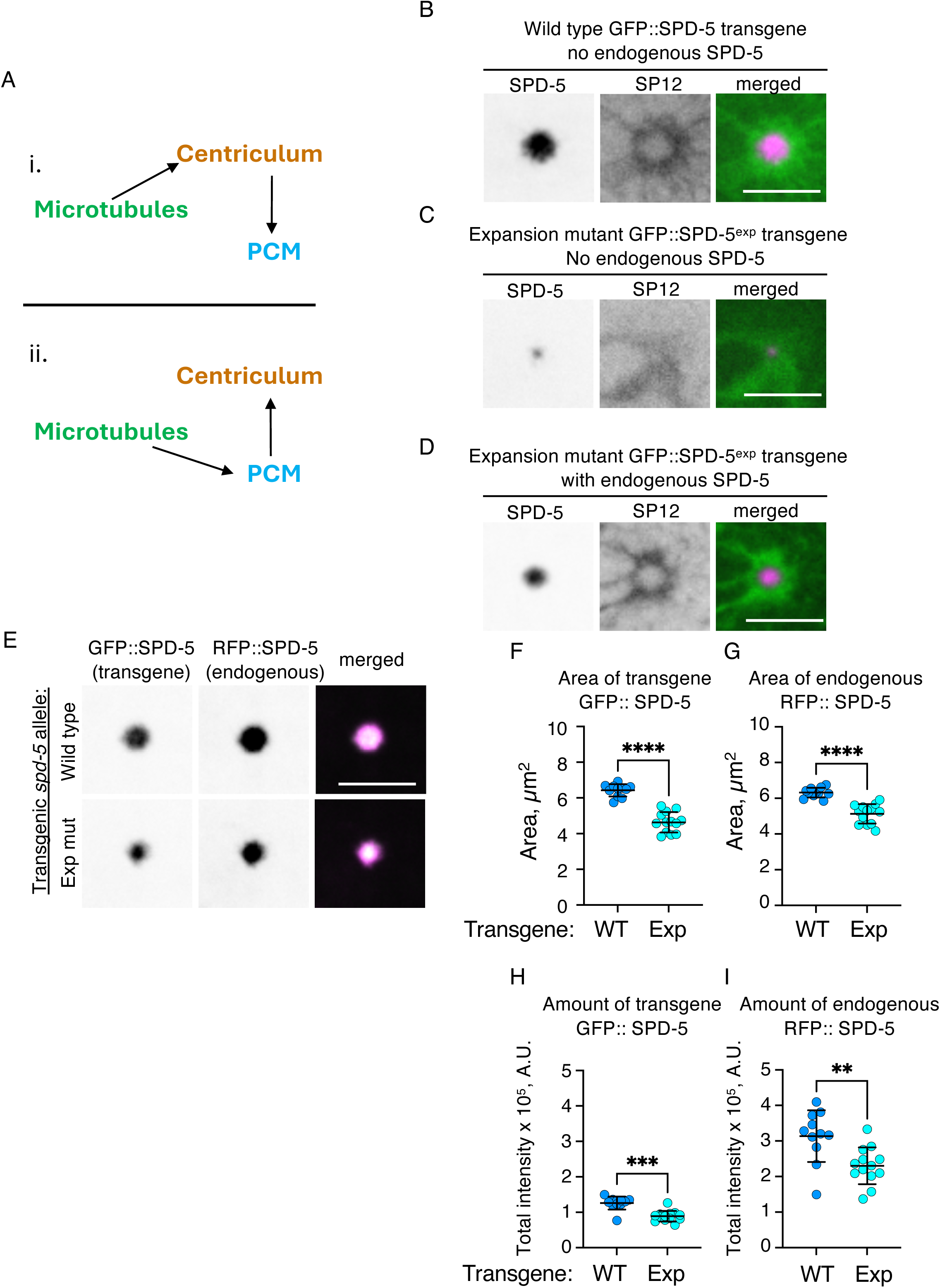
*spd-5* expansion mutant inhibits PCM expansion in the presence of wild type *spd-5*. **A)** Diagram depicting possible relationships between microtubules, the centriculum, and the PCM. See text for more detail. **B, C and D**) Representative images of centricula and PCM from 1-cell embryos at metaphase expressing mCherry::SP12 (green in merged images) and either (B) a *GFP::spd-5* transgene in the absence of endogenous protein (OCF187), (C) *GFP::spd-5^exp^* in the absence of endogenous protein (OCF189), or (D) *GFP::spd-5^exp^* in the absence of presence of endogenous protein (OCF189). The transgenic SPD-5 is shown in magenta in the merged images. In panels B and C, the endogenous gene was down-regulated using RNAi against *spd-5*; transgenes are codon modified such that they are insensitive to the RNAi treatment (Ohta *et al*., 2021). Additional examples of images of embryos such as in panels C and D are shown in Fig S2A and B) Scale bar=5 μm. **E)** Representative images of PCM from 1-cell embryos at metaphase expressing endogenous RFP::SPD-5, and either transgenic GFP::SPD-5 wildtype (OCF201) or transgenic GFP::SPD-5^exp^ (OCF200). Scale bar= 5 μm. **F and G)** Quantification of areas occupied by endogenous RFP::SPD-5 and either transgenic GFP::SPD-5 (panel F) or GFP::SPD-5^exp^ (panel G) from images such as shown in panel E. n=11 and 13 for transgenic wild type and expansion mutant, respectively. p<0.0001 using unpaired t test. Error bars indicate mean and standard deviation. **H and I)** Quantification of protein amount (as reflected by total intensity) occupied by endogenous RFP::SPD-5 and either transgenic GFP::SPD-5 (panel H) or GFP::SPD-5^exp^ (panel I) in the areas quantified in panels F and G. p=0.0004 (panel H) and 0.0034 (panel I), using Mann Whitney test.

SPD-5 has been shown to have two separable functions: (1) centrosome maturation and (2) microtubule nucleation via γ-tubulin recruitment (Ohta *et al*., 2021). Both functions are regulated by the PLK-1 kinase, which phosphorylates different residues on SPD-5. Ohta et al created an RNAi-resistant *spd-5* phospho-mutant allele, *spd-5^exp^*, carrying S653A and S658A substitutions, which they expressed as a transgene in the presence of the endogenous wild type, RNAi sensitive, *spd-5*. As a control, wild type *spd-5* was expressed as an RNAi-resistant transgene. As expected, when SPD-5^exp^ was expressed as the sole SPD-5 version (i.e. endogenous *spd-5* was down-regulated by RNAi), the expansion of the PCM was greatly attenuated compared to a strain expressing transgenic wild type SPD-5 as its sole source of SPD-5 ((Ohta *et al*., 2021) and Figs. 3B and C). However, when transgenic SPD-5^exp^ was expressed in the presence of the endogenous SPD-5 (which is untagged), transgenic SPD-5^exp^ occupied a greater area than without the endogenous protein (compare transgenic GFP::SPD-5^exp^ in Fig. 3C (no endogenous SPD-5) with Fig. 3D (with endogenous SPD-5); see also Fig. S2A and B for more examples). This suggests that SPD-5^exp^ integrates into the wild type SPD-5 lattice despite lacking two PLK-1 phosphorylation sites that contribute to PCM expansion. This is consistent with in vitro findings showing that additional SPD-5 incorporates throughout an existing SPD-5 lattice (Woodruff *et al*., 2017) and with findings by Wueseke et al, who made similar observation with a *spd-5* allele lacking four PLK-1 phosphorylation sites, including the two in SPD-5^exp^ (Wueseke *et al*., 2016). As in Wueseke et al, we also observed a good overlap between the endogenous SPD-5 (fused to tagRFP) and transgenic SPD-5 or SPD-5^exp^ proteins (both fused to GFP) (Fig. 3E), suggesting that both transgenes are integrated throughout the SPD-5 lattice.

Interestingly, the expression of the SPD-5^exp^ transgene in an otherwise wild type background led to a smaller PCM, as judged by the areas occupied by tagRFP-tagged endogenous SPD-5 (Figs. 3E-G). Furthermore, the amounts of both endogenous and transgene SPD-5 at the centrosome were reduced in the presence of the *spd-5^exp^* mutant (Figs. 3 H and I). Thus, SPD-5^exp^ may act as a dominant negative, impeding the ability of the SPD-5 lattice to fully expand. Alternatively, or in addition, the presence of SPD-5^exp^ could have led to reduced overall levels of wild type SPD-5, as proposed by Wueseke et al (Wueseke *et al*., 2016), resulting in a smaller PCM. Note, however, that the fluorescently tagged endogenous SPD-5 may exhibit a genetic interaction with the expansion allele (see below).

### Centriculum size is determined by peri-centrosomal microtubules, not the PCM

Having a condition that reduced the size of the PCM, we next asked whether and how the presence of the *spd-5^exp^* mutant affected the area occupied by peri-centrosomal microtubules. This was done using strains expressing wild type or expansion mutant *spd-5* transgene fused to GFP in the presence of endogenous (untagged) SPD-5, and α-tubulin fused to tagRFP (*tba-1::tagRFP*-T; Fig. 4A). Despite having a smaller PCM in the presence of *spd-5^exp^,* as shown before (Fig. 4B), the amount of tubulin withing the peri-centrosomal region was comparable in the presence of either transgene (Fig. 4C). Moreover, the area occupied by peri-centrosomal microtubules did not decrease due the presence of the expansion mutant transgene (Fig. 4D). As before (Fig. 1E and Fig S1B), peri-centrosomal microtubules occupied a greater area than SPD-5. This area differential increased in the presence of the *spd-5^exp^* mutant (Fig. 4E) due to the smaller area occupied by SPD-5 and an unchanged area occupied by peri-centrosomal microtubules. Thus, peri-centrosomal microtubules were less affected by the presence of the SPD-5^exp^ variant compared to the PCM.

**Figure 4:**
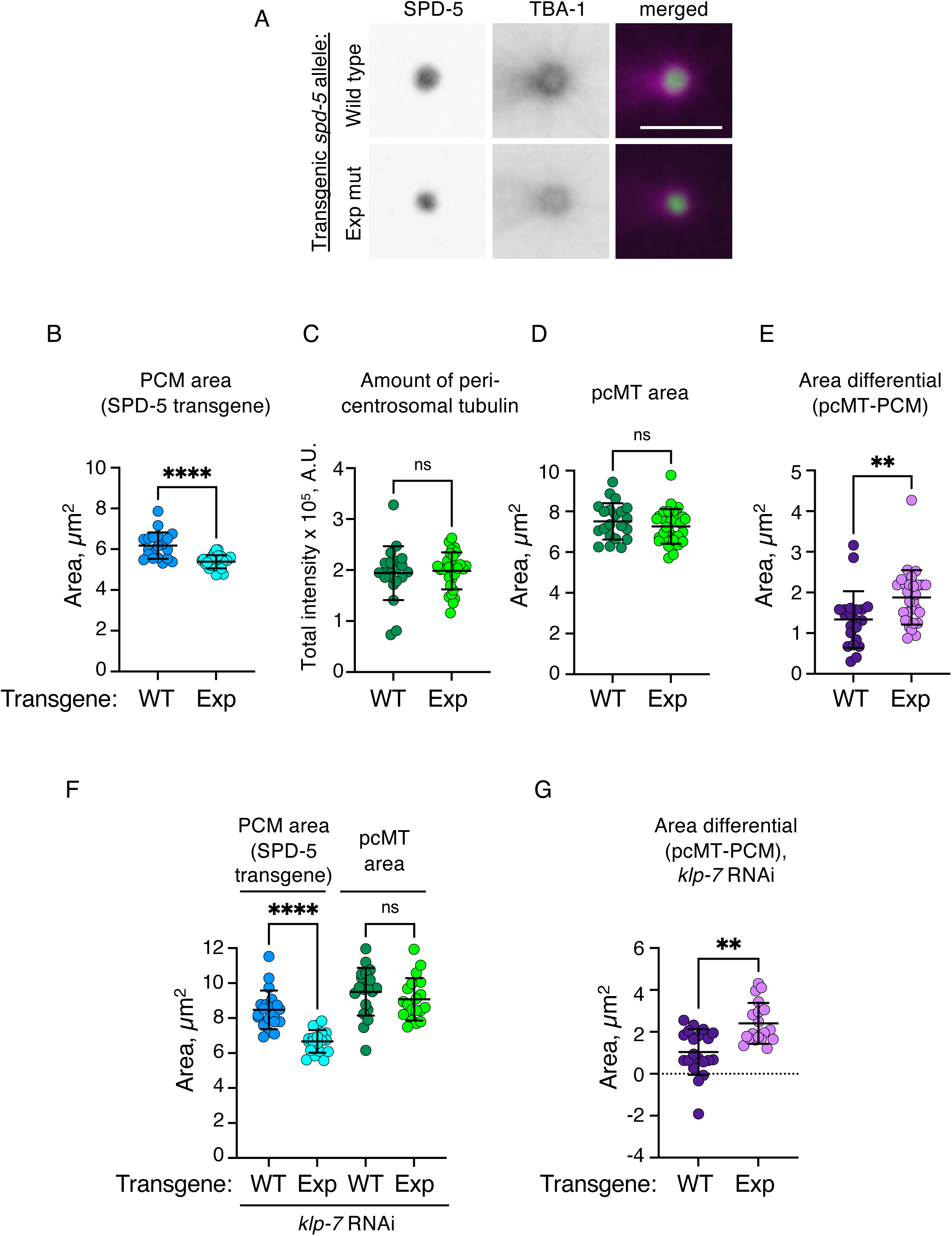
Microtubule nucleation remains unaffected in the presence of SPD-5 expansion mutant. **A)** Representative images of peri-centrosomal microtubules and PCM from 1-cell embryos at metaphase expressing endogenous *tba-1::RFP* and either transgenic *GFP::spd-5* (OCF212) or transgenic *GFP::spd-5^exp^*(OCF213), after control RNAi treatments. Scale bar=5 µm. **B, C and D)** Quantification of PCM area (panel B), amount of peri-centrosomal tubulin (panel C) and the area occupied by peri-centrosomal microtubules (pcMT; panel D) from control RNAi treated embryos such as shown in panel A. n=21 and n=29 for cells expressing transgenic wild type (WT) or expansion mutant (Exp) *spd-5*, respectively. p<0.0001 (panel B) and p=0.3192 (panel D) using unpaired t test, and p=0.6400 (panel C) using Mann Whitney test. Error bars indicate mean and standard deviation. **E)** Quantification of area differential (pcMT area minus PCM area) derived from panels B and D. p=0.0033 using Mann Whitney test. **F)** Quantification of PCM area and pcMT area from the *klp-7* RNAi treated embryos using the same strains as in panels A-E (OCF212 and OCF213). n=20 and n=19 for strains expressing wild type or *spd-5^exp^* transgenes, respectively. p<0.0001 for PCM areas and p= 0.4056 for pcMT areas, using ordinary one-way ANOVA. Error bars indicate mean and standard deviation. **G)** Quantification of area differential (pcMT- PCM, as in panel E) derived from pcMT and PCM areas (panel F) in *klp-7* RNAi treated embryos. p=0.0014 using the Mann Whitney test.

This differential change in size of the PCM and peri-centrosomal microtubules due to *spd-5^exp^* allowed us to examine whether centriculum size follows the PCM or microtubules. Specifically, by decreasing PCM size, the *spd-5^exp^* allele increased the distance (Δ) between the perimeter of the PCM and that of the peri-centrosomal microtubules (Fig. 5Ai). We reasoned that measuring the effect of the *spd-5^exp^* allele on the centriculum void area (determined by the inner perimeter of the centriculum) would allow us to determine if centriculum size tracks with the PCM or the peri-centrosomal microtubules (Fig. 5Aii). To do so, we used the same pairwise combinations used to determine the spatial relationship between the centriculum and the PCM or peri-centrosomal microtubules (Fig. 1B-F), this time in the presence of the wild type or *spd-5^exp^* mutant transgene.

**Figure 5:**
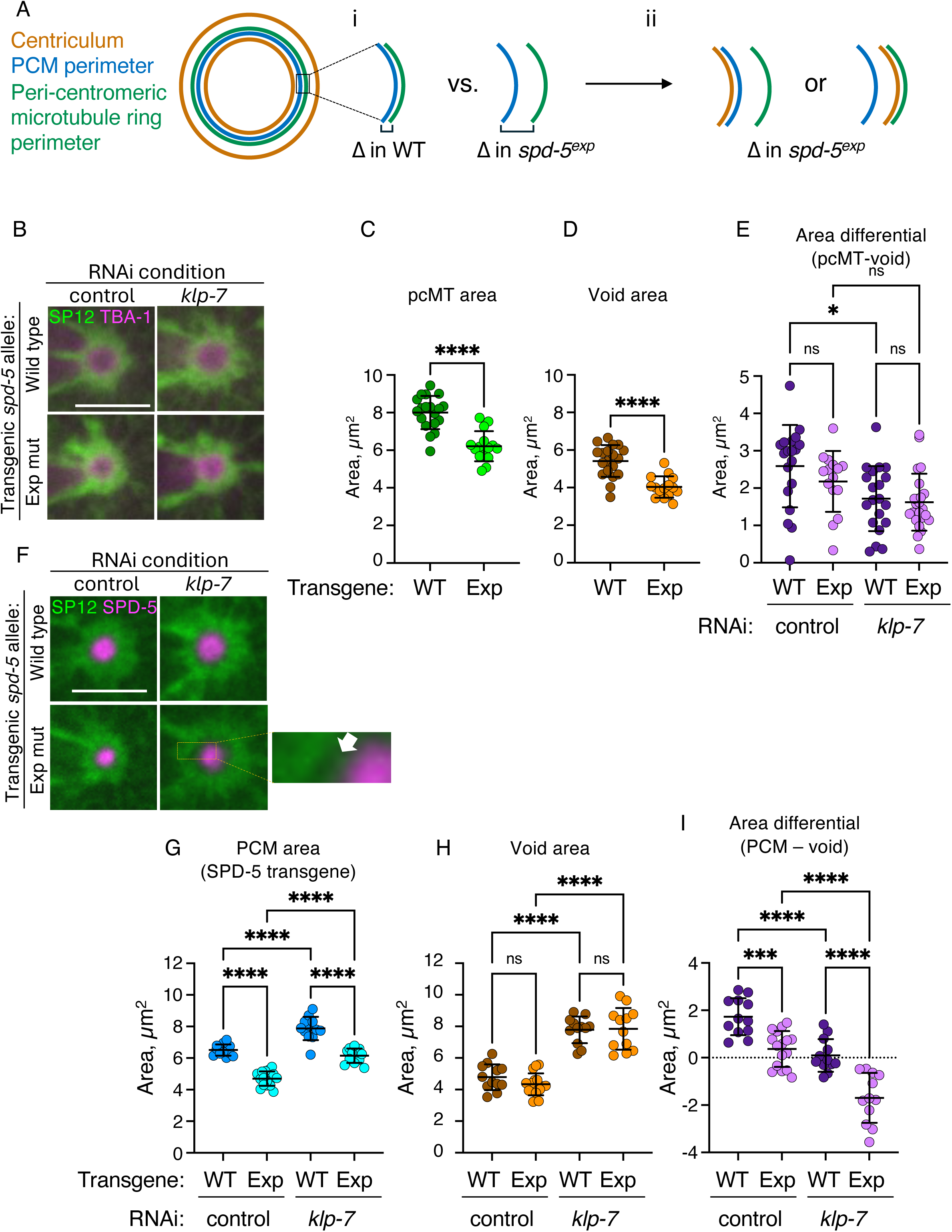
Centriculum size correlates with the position of peri-centrosomal microtubules, not the PCM. **A)** Schematic representation of the spatial relationship between the perimeters of the centriculum (orange), PCM (blue) and the peri-centrosomal microtubules (green). (i) In the presence of *spd-5^exp^*mutant, the area of the PCM, but not the peri-centrosomal microtubules, decreases, and thus the distance between the edges of the two structures should increase. (ii) When considering how the centriculum size, and specifically the area bound but the centriculum’s inner perimeter (the void area), is affected by the presence of *spd-5^exp^*mutant, two possibilities emerge: the edge of the void area follows the PCM (left) or the peri-centrosomal microtubules (right). See text for more detail. **B)** Example of centricula and peri-centrosomal microtubules from 1-cell embryos expressing GFP::SP12 (green in merged image) and TBA-1::RFP (magenta in merged image) and either wild type (OCF214) or *spd-5^exp^* (OCF215) transgenes, following control or *klp-7* RNAi treatments. Scale bar=5 µm. **C and D)** Quantification of pcMT areas (panel C) and void areas (panel D) from control RNAi treated embryos as in panel B. n=19 and n=15 for transgenic wild type and spd-5^exp^ expressing strain, respectively. p<0.0001 for pcMT area comparison using Mann Whitney test, and p<0.0001 for void area comparison using unpaired t test. Error bars indicate mean and standard deviation. **E)** Quantification of area differential (pcMT area minus void area) derived from area values shown in panels C and D and Fig S3C. n=19, 15, 20, 23 for control RNAi, wild type and *spd-5^exp^* transgenes and *klp-7* RNAi, wild type and *spd-5^exp^* transgenes. p-values, from left to right, were 0.5653, 0.013, 0.2353 and 0.9945, using ordinary one-way ANOVA. Error bars indicate mean and standard deviation. **F)** Representative images of centricula and PCM from 1-cell embryos at metaphase expressing *mCherry::SP12* (green in merged images) and either *GFP::spd-5* (OCF187) or *GFP::spd-5^exp^* (OCF189) transgenes (magenta in merged images), following control or *klp-7* RNAi treatment. White arrow in enlarged image points to a gap between the centriculum and SPD-5. Scale bar=5 µm. **G and H)** Quantification of PCM area (panel G) and void area (panel H) from 1-cell metaphase embryos as shown in panel F, following control or *klp-7* RNAi treatment. n=12 all conditions except n=16 for *spd-5^exp^* transgene following control RNAi. For PCM area, p<0.0001 for all pairwise comparisons as determined by ordinary one-way ANOVA. For void area, p-values, from left to right, are 0.6133, <0.0001, <0.0001 and 0.9999, as determined by ordinary one-way ANOVA. Error bars indicate mean and standard deviation. **I)** Quantification of the area differential (PCM area minus void area) derived from area values shown in panels G and H. p-values, as determined by ordinary one-way ANOVA, were <0.0001, except for WT vs Exp, control RNAi, where the p-value was p=0.0003. Error bars indicate mean and standard deviation.

In strains expressing fluorescently tagged SP12 and TBA-1, the *spd-5^exp^* mutant led to a decrease in the area occupied by peri-centrosomal microtubules (Figs. 5B and C) but not in the amount of peri-centrosomal tubulin fluorescence (Fig. S3A). This decrease in peri-centrosomal microtubules area was unlike the situation in strains expressing fluorescently tagged SPD-5 and TBA-1, where *spd-5^exp^*did not affect peri-centrosomal microtubule area (Figs. 4B and C). The reason for this difference is not known. Nonetheless, *spd-5^exp^*also led to a comparable decrease in centriculum size, as seen by the decrease in void area (Fig. 5D) and the area encompassed by the centriculum outer perimeter, COPA (Fig. S3B). Importantly, the reduction in microtubule and void areas were the same (Fig. 5E, control RNAi). In contrast, when we examined the effect of the *spd-5^exp^* transgene on the centriculum in relation to the PCM using strains expressing mCherry-tagged SP12 and GFP-tagged wild type or *spd-5^exp^* transgenes, the effect of *spd-5^exp^*on the PCM was significantly greater than on the centriculum: in these strains, *spd-5^exp^* caused the area of the PCM, but not the centriculum, to shrink (Figs. 5F-I, control RNAi). In fact, in several cases the size of the PCM was smaller than that of the centriculum void (see negative values in Fig. 5I). Taken together, these data suggest that centriculum size follows the behaviors of the peri-centrosomal microtubules rather than the PCM.

To further examine the dependence of centriculum size on microtubules vs. the PCM, we stabilized microtubules in strains expressing wild type or *spd-5^exp^* transgene using RNAi against *klp-7*. We reasoned that if centriculum size is determined by microtubules, stabilizing microtubules in the presence of the *spd-5^exp^* transgene might further increase the distance between the PCM and the inner perimeter of the centriculum. As expected, (Maheshwari *et al*., 2023), *klp-7* RNAi led to expansion of the centriculum, PCM and peri-centrosomal microtubule area (for PCM and peri-centrosomal microtubules areas, compare the values in Fig. 4F (*klp-7* RNAi) with Fig. 4B and 4D (control RNAi). The effect of *klp-7* RNAi on the PCM in a different strain is shown in Fig. 5G; for the centriculum void area and COPA, see Fig. 5H and Fig. S3B). Following *klp-7* RNAi treatment, the effect of *spd-5^exp^*was again greater on the PCM than on the pericentrosomal microtubules (Figs. 4F and G), while area differential between the centriculum void area and the peri-centrosomal microtubules remained the same (Fig. 5E; pcMT and void areas in the presence of *klp-7* RNAi are shown in Fig. S3C). However, when strains expressing the transgenic wild type or *spd-5^exp^* mutant and mCherry::SP12 were treated with RNAi against *klp-7*, a visible “gap” appeared between the PCM and the inner perimeter of the centriculum (Fig. 5F, see enlarged area). While *klp-7* RNAi led to an overall increase in PCM area, the presence of the *spd-5^exp^* allele led to reduced PCM area (Fig. 5G) without affecting the centriculum void area (Fig. 5H). Consequently, stabilizing microtubules through *klp-7* RNAi exacerbated the difference between the void area and the PCM such that in all cases, the PCM was smaller than the inner perimeter of the centriculum (Fig. 5I, note negative values in the presence of the *spd-5^exp^*allele following *klp-7* RNAi). In fact, *klp-7* RNAi alone was sufficient to increase the area differential between the void area and the PCM (Fig. 5I). Taken together, these data suggest that the centriculum behaves the same as microtubules, not the PCM, and that centriculum size is determined by microtubules.

In the experiments described above, the size of the PCM was assessed by the fluorescently tagged transgenes. While there was good agreement between the areas occupied by the transgenic and the endogenous SPD-5s (Fig. 3F and G), we attempted to examine the spatial relationship between the centriculum and the PCM using an endogenously tagged SPD-5. Unexpectedly, the *spd-5^exp^* transgene and the endogenously tagged *spd-5* exhibited a genetic interaction, resulting in abnormal distribution of both proteins (Figs. 6A and Fig. S4), reminiscent of SPD-2 distribution (see for example (Pelletier *et al*, 2004)). We assume that when the endogenous *spd-5* was not tagged, its distribution was uniform, as judged by the uniform distribution of the GFP::SPD-5^exp^ transgene, which on its own cannot expand (Fig. 3C and D and Fig. S4A). However, when the endogenous *spd-5* was fused to a fluorescent protein, its distribution in the presence of the transgenic *spd-5^exp^* mutant becomes abnormal, with a high concentration in the center of the centrosome, and a much lower concentration in the majority of the PCM (Fig. S4B and C). Under these conditions, the distribution of the GFP::SPD-5^exp^ transgene was similarly abnormal (Fig. S4B), in further support of the similar distributions of the endogenous and transgene proteins. Despite this abnormal distribution, we were still able to determine the area occupied by endogenously expressed SPD-5, including the fainter periphery, using the same thresholding method applied to all other area measurements (Fig. S4D). As before, the presence of the *spd-5^exp^* mutant transgene decreased the amount of SPD-5 at the centrosome, and this pattern was unaffected by down-regulating *klp-7* (Fig. 6B). Also as before, down-regulation of *klp-7* increased PCM area, and the presence of the *spd-5^exp^* mutant decreased PCM areas relative to the wild type transgene in both control and *klp-7* RNAi conditions (Fig. 6C). Consistent with the observations above, the presence of the *spd-5^exp^* mutant did not reduce the size of the centriculum void volume, and down-regulation of *klp-7* even increased it (Fig. 6D). Consequently, the area differential between the PCM and the centriculum increased in the presence of the *spd-5^exp^* mutant, and this difference was further exacerbated when *klp-7* was down-regulated (Fig. 6E).

**Figure 6:**
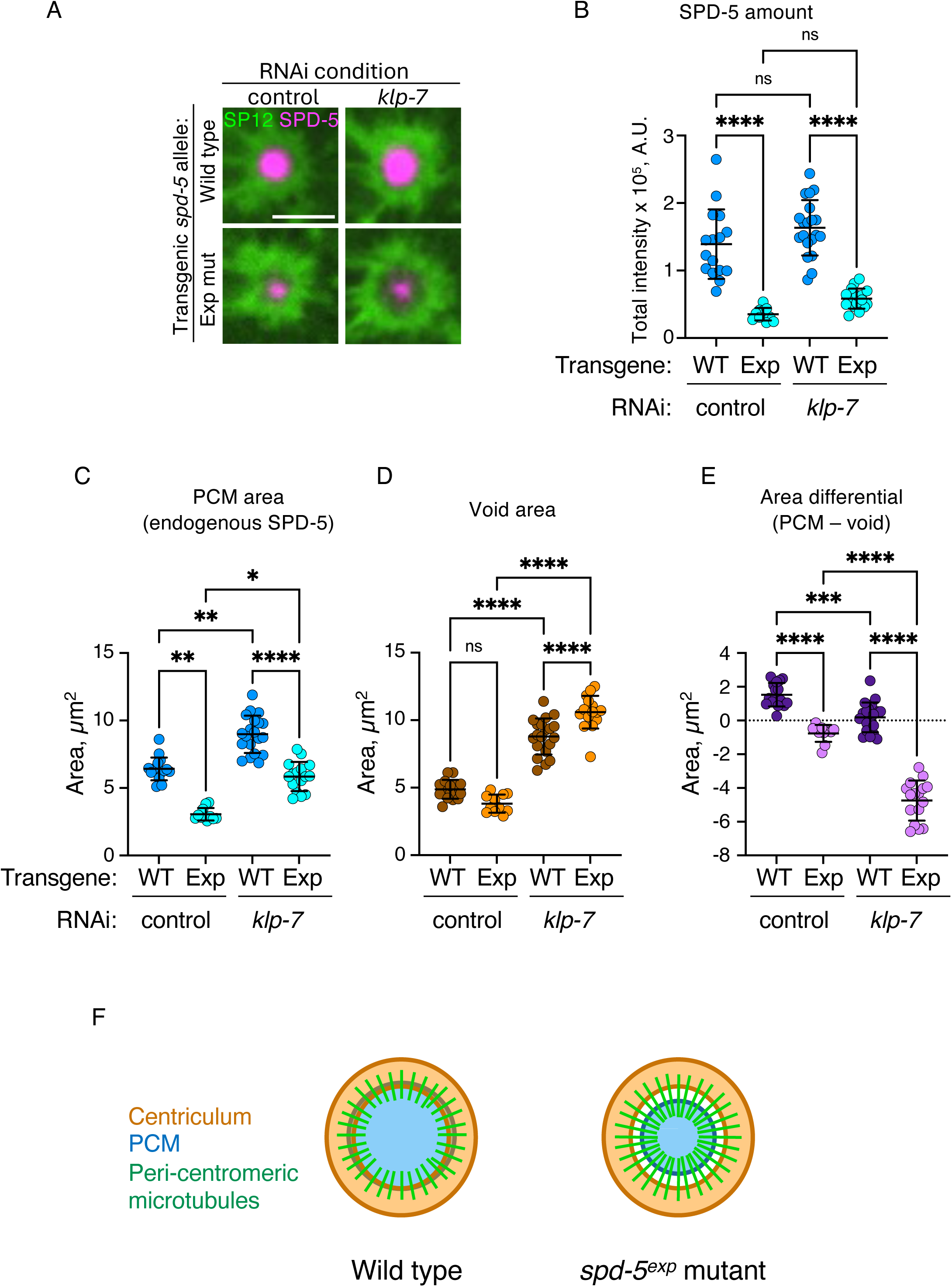
Spatial relationship between the centriculum and the PCM using an endogenously tagged *spd-5* in the presence of *spd-5^exp^* mutant. **A)** Representative images of centricula and PCM from 1-cell embryos at metaphase expressing RFP-tagged endogenous *spd-5* (RFP::SPD-5, in magenta) and GFP::SP12 (in green) in strains expressing either wild type (OCF234) or *spd-5^exp^* (OCF233) transgenes, following control or *klp-7* RNAi treatment. Scale bar=5 µm. **B, C and D)** Quantification of SPD-5 amount, PCM areas and void areas from 1-cell metaphase embryos treated as shown in panel A. n=16 and 11 for wild type and *spd-5^exp^* transgenes treated with control RNAi, and n=20 and n=16 for wild type and *spd-5^exp^*transgenes treated with *klp-7* RNAi. p-values, from left to right, were as follows: panel B: <0.0001, 0.1795, 0.3578 and <0.0001, as determined by ordinary one-way ANOVA; panel C: 0.0018, 0.0023, 0.0286 and <0.0001, as determined by Kruskal-Wallis test; panel D: 0.0547, and the rest were <0.0001, as determined by ordinary one way ANOVA. Error bars indicate mean and standard deviation. **E)** Quantification of the area differential (PCM area minus void are) derived from data shown in panels C and D. p-values, as determined by ordinary one-way ANOVA, were <0.0001 except for p=0.0001 for wild type, control vs *klp-7* RNAi. Error bars indicate mean and standard deviation. **F)** Diagram illustrating the effect of the spd-5^exp^ transgene on the centriculum (orange), peri-centrosomal microtubules (green) and PCM (blue). See text for more detail.

Taken together, these data suggest two things (Fig. 6F): First, that centriculum size is likely determined by sum forces exerted on it by peri-centrosomal microtubules. Second, because the area occupied by pericentrosomal microtubules change little or none when the PCM “shrinks” due to the presence of the *spd-5^exp^*mutant, and because microtubules nucleate from the surface of the PCM, our data suggest that the length of pericentrosomal microtubules is not an inherent property of these microtubules, because their length, from the surface of the PCM to the inner perimeter of the centriculum, must be different in wild type vs *spd-5^exp^* expressing cells (Fig. 6F). Rather, our data suggest that the majority of pericentrosomal microtubules extend from the surface of the PCM until they encounter the centriculum, where most of them terminate due to obstruction by the centriculum membrane; only a small fraction can extend past the centriculum to form astral or spindle microtubules. As such, the centriculum may serve as a microtubule filter, a possibility that is explored next.

### Centriculum porosity correlates with the ability of microtubules to traverse the centriculum

As noted above, two types of centrosome-nucleated microtubules extend beyond the centriculum: astral microtubules, which extend towards the cell cortex, and spindle microtubules, which extend towards chromosomes. The density of spindle microtubules is much greater than astral microtubules (Fig. 7A, metaphase). Spindle microtubules are only evident in metaphase; prior to that, only astral microtubules can be observed outside the centriculum, and no microtubules have entered the pronuclei (Fig. 7A, prophase). If the centriculum serves as a microtubule filter, we would expect that at metaphase, the centriculum would be more porous on the side facing the chromosomes vs. the side facing the cortex. To examine this possibility, we measured the effective pore size that is available for microtubules nucleated at centrosomes to pass through the centriculum using volume electron microscopy data acquired via focused ion beam-scanning electron microcopy (FIB-SEM; (Maheshwari *et al*., 2023)).

**Figure 7:**
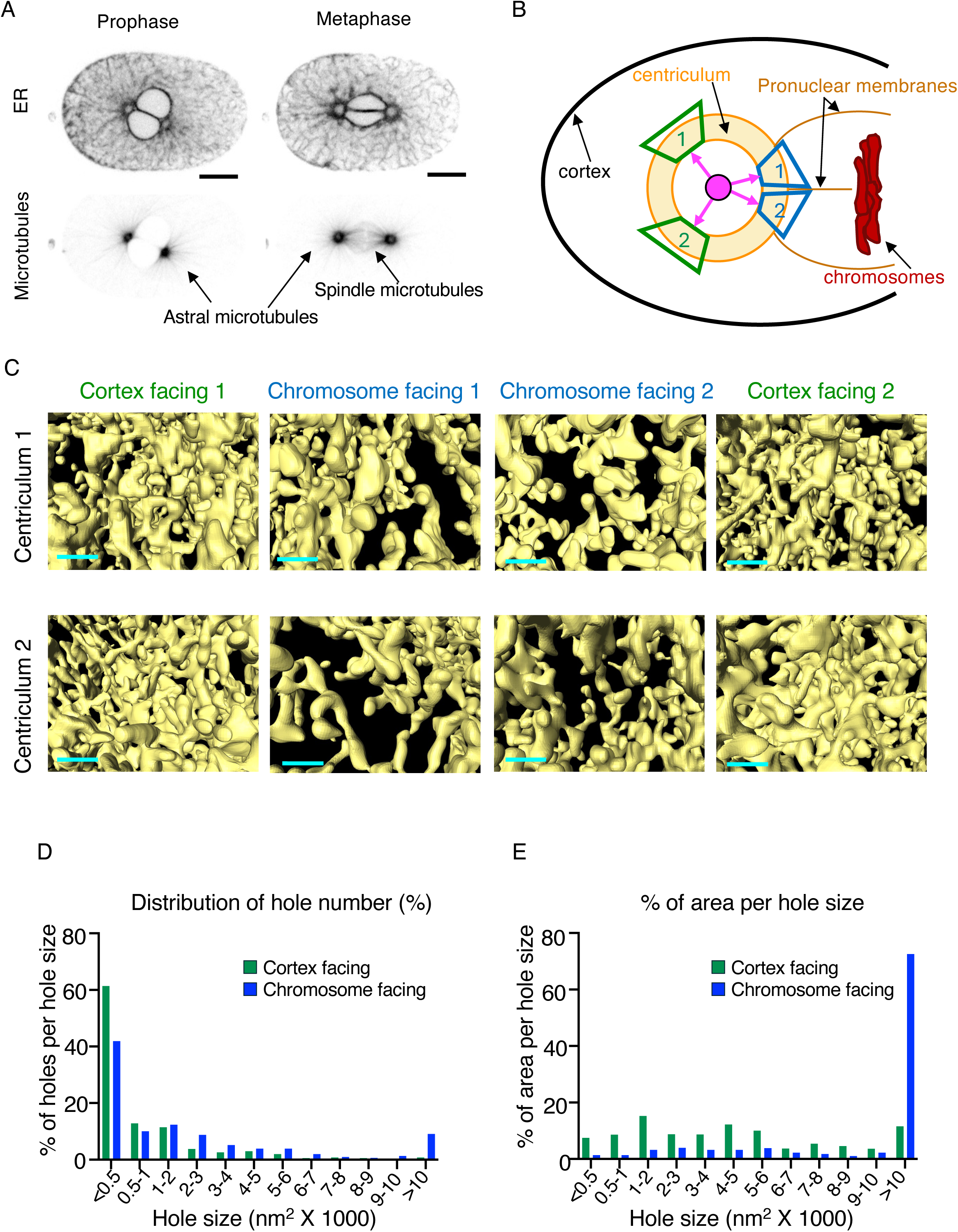
Porousity of the centriculum facing the cortex. **A)** Representative images of 1-cell *C. elegans* embryos expressing mCherry::SP12 to visualize the centriculum and GFP::TBA-2 (α-tubulin) to visualize microtubules (strain OCF181) at prophase (left side) and metaphase (right side). Note that at prophase, microtubules are excluded from the nucleoplasm, and that in metaphase, the density of spindle microtubules is greater than astral microtubules. Scale bar=10 μm. **B)** Diagram depicting the centriculum (in orange) fused to the two pronuclei (brown lines) at a metaphase 1-cell embryo. Chromosomes (in red) and the cortex (black line) are also indicated. The areas analyzed for pore size are shown as blue (chromosome facing) and green (cortex facing) trapezoids. A central fiducial (in pink) that is equidistant (pink arrows) to the areas analyzed is also shown. **C)** Cortical and pronuclear centriculum segments based on FIB-SEM data (Maheshwari *et al*., 2023) from two centricula, shown side by side. The images were taken from the central fiducial. Scale bar=200 nm. **D)** Binned frequency distribution of holes present on the cortical side (in green) and chromosome side (in blue) of centricula in 1-cell metaphase embryos. Bin size range, as hole area, is shown on the x-axis. n=308 and 507 for chromosomal and cortex side holes, respectively, each containing data pooled from 6 images taken from 3 centricula (2 images per centriculum). **E)** Binned frequency distribution of the percentage of total open area per hole size range, for holes on the cortical side (green) and the pronuclear side (blue) using the same data as in panel D.

We created a fiducial in the center of segmented metaphase centricula, and imaged a portion of the centriculum, from that fiducial, against a black background (Fig. 7B and C; images were taken from within the centrosome towards either the cortex or the chromosomes). Qualitatively, it already appeared that the holes on the chromosomal side are larger than those on the cortical side (compare the two central panels, chromosome-facing 1 and 2, to the side panels, cortex-facing 1 and 2). To maximize having the openings imaged *en face*, the angles between the center line and the top or bottom of any image were kept to a minimum (for cortex facing areas: 23.8° ± 2.82°, average and standard deviation, n=6; for chromosome facing areas: 22.8° ± 2.23°, average and standard deviation, n=6). We then determined the area of each opening, defined as “hole”, that can be seen through the centriculum. Considering that the diameter of a microtubule is ∼25 nm, the area of the smallest hole through which a microtubule can pass would be 490 nm^2^ (𝜋 x 12.5^2^) and probably larger since most holes are not perfect circles. The fraction of cortex-facing centriculum area that was open, namely the sum of all hole areas out of the total area imaged, was 7.85 ± 1.66% (n=6, from 3 centricula), while on the chromosome side it was significantly greater: 20.66 ± 3.75%, (n=6 from 3 centricula; p=0.009 as determined by chi square test), consistent with the chromosome-facing side of the centriculum having larger opening. In the case of cortex-facing holes, the cross sections of 61.34% of the holes, accounting for 7.5% of the total open area, were below 500 nm^2^, likely too small to allow for a microtubule to pass (Fig. 7D and E). The percent of chromosome facing holes that had a cross section below 500 nm^2^ was 41.88%, and this accounted for only 1.4% of the total open area on the chromosomal side (Fig. 7D and E). On the other side of the hole-size spectrum, 11.56% of the open area facing the cortex were from holes that were larger than 10,000 nm^2^ (0.8% of all holes), compared to 74.52% on the chromosome-facing side (9.1% of all holes) (Fig. 7E). Thus, the centriculum side facing the chromosomes, through which spindle microtubules pass, is significantly more porous than the side facing the cortex, where astral microtubules pass. Whether the different degrees of porosity are caused by centriculum geometry or by properties of spindle microtubules remains to be determined (see Discussion).

Given that only a small fraction of the centriculum area facing the cortex is available for the microtubules to go through, and the fact that the thickness of centriculum at metaphase is around 1 µm (Maheshwari *et al*., 2023), we predicted that the microtubules that become fully elongated astral microtubules are those that take the shortest path through the centriculum, and are thus less likely to encounter centriculum membrane. To test this hypothesis, we used ER tomography datasets of 1-cell embryos at metaphase (Maheshwari *et al*., 2023; Redemann *et al*., 2017) and measured the angle of microtubules as they pass the inner perimeter of the centriculum relative to the radial vector, the line that is perpendicular to the tangent line (Fig. 8A). The shortest path through the centriculum would be that of the radial vector (angle= 0°); increasing angles between a microtubule path and the radial vector would mean longer paths through the centriculum. We then determined whether the microtubule passed through the centriculum (denoted as “continuing”) or whether the microtubule terminated within the centriculum (denoted “stopped”). On the cortex-facing side, we measured angles for 113 microtubules, of which 21 continued (18.6%), and on the chromosome-facing side we measured angles for 146 microtubules, of which 40 continued (27.4%). In both cases, the distribution of angles of microtubules that passed all the way through the centriculum was significantly narrower, and closer to the radial vector, than the range of angles for microtubules that ended within the centriculum (Fig. 8B). Nonetheless, our data suggest that the centriculum serves as a passive microtubule filter, blocking most microtubules from extending past the centriculum.

**Figure 8:**
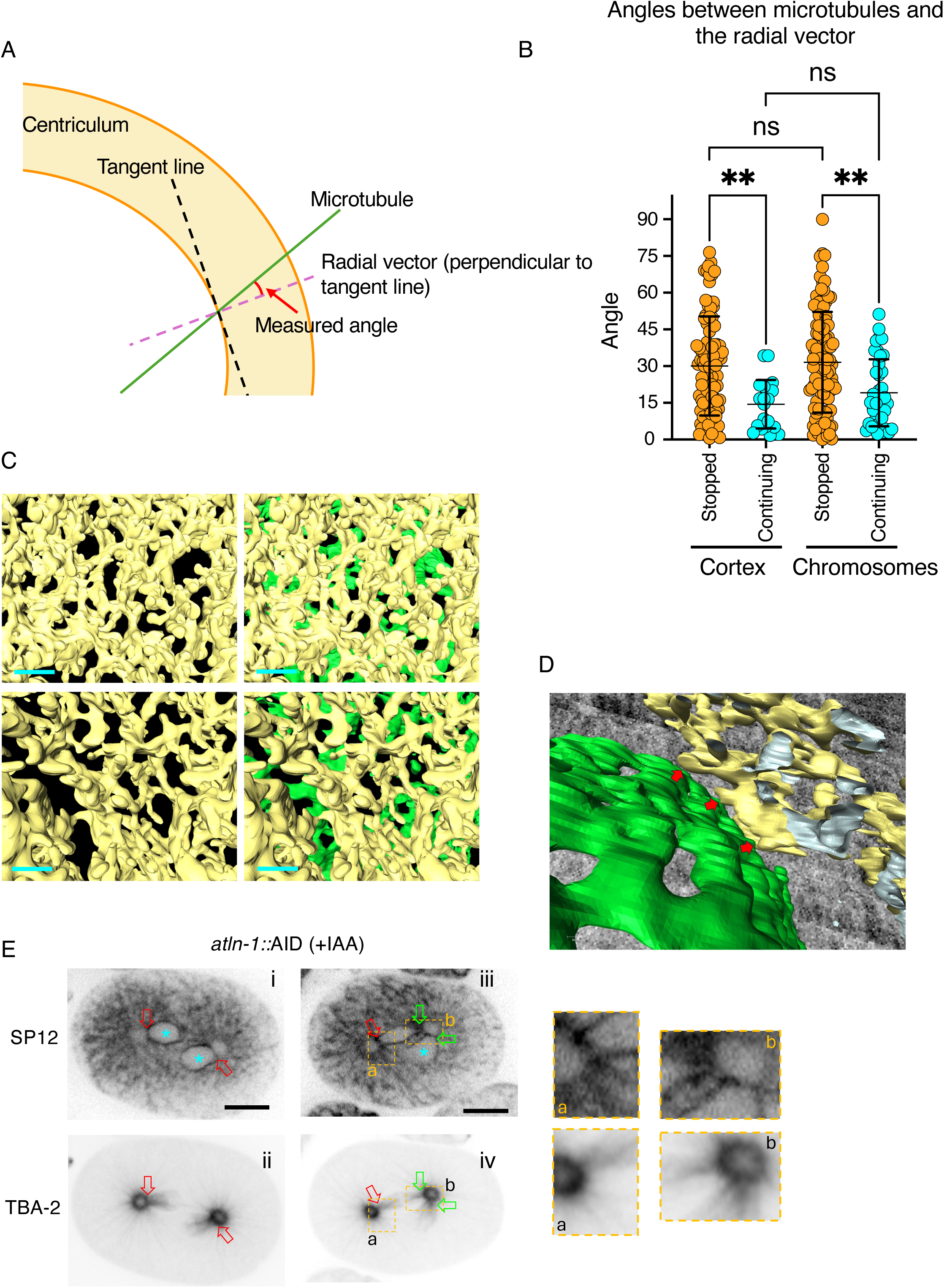
**A)** Diagram depicting the angle of a microtubule (green line) passing through the centriculum (in orange) relative to the radial vector (dashed purple line). The actual measurement was done between the microtubule and the tangent line (dashed black line) and the value obtained was subtracted from 90 to obtain the angle to the radial vector, which is perpendicular to the tangent line. **B)** Quantification of angles between of stopped (in orange) and continuing (in cyan) microtubule and radial vector for astral (cortex-facing) and spindle (chromosome-facing) microtubules. Based on EM tomography data (Redemann *et al*., 2017). p=0.0029 (for cortex-facing) and 0.0018 (chromosome-facing) using one-way ANOVA with multiple comparisons. Error bars indicate mean and standard deviation. **C)** Left panel: Representative images of a segmented area based on FIB-SEM data (Maheshwari *et al*., 2023) from the pronuclear side of two prometaphase centricula (in yellow), visualized from the center of the centrosome. Right panel: the same images with segmentation of nuclear membrane remnants (in green) that are between the centriculum and the prometaphase chromosomes (not shown). Scale bar=200 nm. **D)** Areas of possible contact (indicated by red arrows) between a prometaphase centriculum (in yellow) and remnants of the nuclear membrane (in green) based on FIB-SEM data (Maheshwari *et al*., 2023). In this image, the centrosome is to the upper right side while the chromosomes are to the lower left side. **E)** Two 1-cell embryos expressing mCherry::SP12 and GFP::TBA-2 from worms in which atlastin was downregulated using an auxin-inducible degron (AID) system (OCF183, see (Maheshwari *et al*., 2023)). Red arrows point to sites where a centriculum has fused to a single pronucleus. Green arrows point to sites where a centriculum has fused to both pronuclei. The same arrows are shown in the bottom panels, where they are adjacent to sites where spindle microtubules extend into the pronuclei. Blue asterisks indicate pronuclei that fuse to only one centriculum. The enlarged areas show sites of centriculum-pronuclear fusion. Note that unlike the wild type situation (Fig 7A), under these conditions the two pronuclei fail to align and do not form extensive pronuclear interfacial contacts. Scale bar=10 μm.

To form a spindle, microtubules must enter the space around the chromosomes. Prior to metaphase, an intact nuclear envelope prevents centrosome-nucleated microtubules from entering the nucleoplasm. Just before metaphase, two events take place: the nuclear envelope loses its integrity specifically next to centrosomes (Hachet *et al*, 2012), and the centriculum fuses with the nuclear membrane (Maheshwari *et al*., 2023). In this regard, the larger hole size of the centriculum on the chromosomal side is unlikely to be the only reason why spindle microtubule density is greater than that of astral microtubules. At prometaphase, the percent of open centriculum area (14.64 ± 4.14%, n=6 from 3 centricula), distribution of hole size, and percent area per hole size (Fig. S5A and B) on the chromosomal side of the centriculum were similar to those of metaphase centricula. Effectviely, however, the number of holes available for microtubules to reach the chromosomes is significantly smaller, because at prometaphase, remnants of the nuclear membrane are still present, creating an additional barrier for microtubules to enter the nucleoplasm (Fig. 8C, note the presence of nuclear membrane (in green) masking many of the openings). At prometaphase the centriculum is adjacent to the nuclear membrane, with a few possible contact sites (see, for example, Fig. 8D), but there are no membrane junctions that indicate actual fusion. In contrast, at metaphase, the centriculum is completely fused with the remnants of the nuclear membrane (Maheshwari *et al*., 2023), possibly in an atlastin-dependent manner ((Araújo *et al*., 2023) and see below). Therefore, at metaphase, there are no other membranes between the centriculum and the chromosomes. Thus, the fusion of the centriculum and mitotic remnants of the nuclear membrane may reduce membrane barriers in the process of spindle formation.

A further indication that centriculum-nuclear membrane fusion is important for spindle microtubule formation is the state of metaphase 1-cell embryos following atlastin down-regulation. Atlastin is a GTPase needed for the fusion of ER tubules (Hu & Rapoport, 2016; Orso *et al*, 2009). We down-regulated atlastin using a degron construct, under conditions that partially lower the amount of atlastin in the embryo (Maheshwari *et al*., 2023). Under these conditions, for unknown reasons, pronuclei have difficulties aligning, and they sometimes fuse with only one of the two centricula (Fig. 8E, see pronuclei marked in blue asterisks in panels i and iii), leading to the formation of monopolar spindles in those pronuclei. Centricula at metaphase are normally fused to both pronuclei, resulting in two dense clusters of spindle microtubules on either side of the membrane between the two pronuclei (for example, see Fig. 7A). However, when *atln-1* is down-regulated, a centriculum may fuse with only one of the two pronuclei (Fig. 8E, red arrows). Under these conditions, dense microtubules can be observed only where the centriculum is fused to the pronuclei, while the rest of the centriculum is surrounded by lower density, astral microtubules (Fig. 8E and enlarged areas from panels iii and iv). This suggests that robust microtubule elongation past the centriculum to form the spindle’s dense microtubule array can only happen once the centriculum fuses with a pronucleus. Taken together, we proposed that formation of high-density microtubules on the chromosome-facing side of the centriculum is facilitated by two processes: the presence of large openings that traverse the centriculum, and the fusion of the centriculum with remnants of the nuclear membrane.

## Discussion

In this study, we tested the hypothesis that the centriculum acts as a partial barrier, or filter, preventing the extension of some, but not all, centrosome nucleated microtubules. Our measurements show that the PCM and peri-centrosomal microtubules either abut or slightly penetrate the inner perimeter of the centriculum (Fig. 1). We further observed, both here and in our previous study, that changing microtubule stability or number affects both PCM size and centriculum size ((Maheshwari *et al*., 2023) and Figs 2, 5G-I, 6 and Figs. S1 C and D, and S3)). This suggested that microtubules affect centriculum size either directly, potentially by exerting force on the centriculum and thus determining its size, or indirectly, by affecting PCM size, which in turn affects centriculum size. To gain insight into the spatial dependencies of these three structures, we used a *spd-5* allele, *spd-5^exp^*, that is defective in PCM expansion (Ohta *et al*., 2021). Expressing both *spd-5^exp^*, as a transgene, and the endogenous wild type *spd-5* resulted in a smaller PCM. An inhibitory effect by SPD-5 mutant proteins on the SPD-5 lattice has been observed previously: Nakajo et al. observed this phenotype with a 272 amino acid C-terminal deletion, which did not encompass the residues mutated in SPD-5^exp^ (Nakajo *et al*., 2022), while Wueseke et al. observed it with a SPD-5 variant lacking four PLK-1 phosphorylation sites, SPD-5^4A^, including the two in SPD-5^exp^ (Wueseke *et al*., 2016). Rios et al found that PLK-1 phosphorylation of SPD-5 reduces intramolecular interactions, allowing these domains to interact intermolecularly (Rios *et al*., 2024). We therefore propose that the SPD-5^exp^ variant acts as a dominant negative by incorporating into the SPD-5 lattice but not contributing fully to intermolecular connections, thereby impeding lattice expansion. Alternatively, or in addition, the presence of transgenic mutant *spd-5* may have led to reduced levels of endogenously expressed *spd-5* (Wueseke *et al*., 2016), thus resulting in a smaller PCM.

Importantly, expressing both transgenic *spd-5^exp^* together with the endogenous wild type *spd-5* resulted in a smaller PCM without reducing the area occupied by the peri-centrosomal microtubules (Fig. 4A-D). The same was true when worms were treated with RNAi against *klp-7*, which stabilizes microtubules: while the areas occupied by the PCM and peri-centrosomal microtubules was overall greater due to *klp-7* down-regulation (Maheshwari *et al*., 2023), only PCM area, but not peri-centrosomal microtubule area, was affected by the presence of SPD-5^exp^ (Fig 4F). In other words, expression of the *spd-5^exp^*allele led to a greater difference in the areas occupied by peri-centrosomal microtubules compared to the PCM (Fig 5E and G). This experimental setup allowed us to address two questions: the first was what determines centriculum size. By examining centriculum size relative to the PCM vs per-centrosomal microtubules in the presence of *spd-5^exp^*, we showed that the centriculum correlates with microtubules, not the PCM (Fig 5). In fact, under *klp-7* RNAi treatment, a gap was visible between the inner perimeter of the centriculum and the PCM (Fig 5F). The increase is distance between the centriculum and the PCM was true regardless of whether PCM area was determined using the transgene or endogenous SPD-5 (Fig 6 E). Thus, our data are consistent with the possibility that centriculum size is determined by microtubules, not by the PCM. This further suggests that PCM size could be affected by the size of the centriculum, such that, for example, when the centriculum is unusually small due to reduced microtubule number or stability, the centriculum could lead to PCM compaction (Fig 2). This is an intriguing possibility, as it suggests that the PCM lattice is malleable, able to expand or fold inwards in response to forces acting upon it, akin to a Hoberman Sphere (https://www.hoberman.com/portfolio/hoberman-sphere-toy/). That the PCM is malleable was also proposed by Laos et al, who noted that the PCM lattice *in vivo* can grow by incorporating SPD-5 anywhere within the lattice (Laos *et al*., 2015). This property could contribute to the PCM’s ability to deform in response to forces exerted on it by astral microtubules (Enos *et al*., 2018), where the PCM could stretch rather than break. This property, known as ductility, was also proposed by Mittasch et al based on the PCM’s deformability during anaphase in response to external forces (Mittasch *et al*., 2020).

The second question this experimental set up allowed us to address is whether peri-centrosomal microtubules are short due to an inherent property. Numerous in vitro studies, using tubulin concentrations that were significantly lower than in the 1-cell *C. elegans* embryo, have examined microtubule length either directly or indirectly, and in none of them were microtubules as short as the *C. elegans* 1-cell peri-centrosomal microtubules (Arnal *et al*, 2004; Belmont *et al*, 1990; Chrétien *et al*, 1995; Farrell *et al*, 1987; Fygenson *et al*, 1994; Gould & Borisy 1977; Grego *et al*, 2001; Kuriyama & Borisy, 1981; Mitchison & Kirschner, 1984; Verde *et al*, 1990), suggesting that something is preventing the extension of peri-centrosomal microtubules. Ideally, we would have liked to remove the centriculum entirely and examine the effect on peri-centrosomal microtubule length. Unfortunately, we have yet to identify a condition that eliminates the centriculum. Nonetheless, the *spd-5^exp^*allele created a condition where limits to peri-centrosomal microtubule length could be examined. Microtubules are nucleated from the outer portion of the PCM. In the presence of the *spd-5^exp^* allele, the PCM is smaller, but the area occupied by peri-centrosomal microtubules, as determined by the outer periphery of this structure, remains unchanged (Fig 4B, D and F). This suggests that in the presence of the *spd-5^exp^*allele, peri-centrosomal microtubules are longer than in the control (Fig 6F). If the length of peri-centrosomal microtubules were a function of an inherent property of these microtubules, the area occupied by these microtubules would have been smaller in the presence of SPD-5^exp^. Thus, we propose that the accumulation of peri-centrosomal microtubules is a consequence of the presence of the centriculum, which blocks the extension of a subset of microtubules that are nucleated by the centrosome. While elongating microtubules have the capacity to distort a membrane sheet (for example, see (Zimmerman & Chang, 2005)), we propose that the reticular nature of the centriculum, creating a membrane lattice, allows it to resist forces exerted on it by microtubules. Consistent with this possibility, the microtubules that manage to pass through the centriculum take, on average, a shorter route than microtubules that stop within the centriculum (Fig 8A and B). If microtubules were agnostic to the presence of the centriculum, there would be no difference in the angle relative to the radial vector of microtubules that terminate within the centriculum and those that extend past it. A similar accumulation of peri-centrosomal microtubules was observed in other organisms that also have peri-centrosomal ER accumulation (Kiyomitsu *et al*., 2024; Rollins & Blankenship, 2023; Xie *et al*., 2025).

While the centriculum may act as a barrier to the extension of peri-centrosomal microtubules, it is still permissive to extension of astral and spindle microtubule extension. The density of microtubules extending toward the chromosomes by far exceeds the density of microtubules extending toward the cortex (Figs. 1A and 7A). This difference in density correlates with the size of centriculum “holes” and the overall open area, which is significantly greater on the side of the centriculum that faces the chromosomes compared to the side facing the cortex (Fig 7). We imagine two non-mutually exclusive reasons why centriculum holes are larger on the side facing the chromosomes: first, prior to metaphase, this side of the centriculum is adjacent to the pronuclear membrane and is not as wide as the parts facing the cytoplasm (Maheshwari *et al*., 2023), perhaps due to a spatial restriction of ER-derived membrane accumulation. If we assume that the centriculum is a passive barrier, the wider it is, the more microtubules it will block from extending. Second, the centriculum is likely a dynamic structure, much like the rest of the ER. At metaphase, microtubules that reach the chromosomes are stabilized, likely restricting the ability of the centriculum to remodel around these microtubules. Under these conditions, it is possible that additional microtubules can pass alongside the stable microtubule, further reducing the ability of the centriculum to remodel at that site and establishing a “hole” whose size may continue to increase. Consistent with this possibility, we observe a high density of microtubules extending past the centriculum only after the centriculum fuses with remnants of the pronuclear membranes (Fig 8E). This explanation may also apply to cortical microtubules, which although not stabilized, may still promote the passage of additional microtubules through the centriculum alongside them. Indeed, the distribution of astral microtubules is not uniform (Fig. 7A), perhaps due to the filter properties of the centriculum.

Taken together, we propose that the centriculum acts as a partial barrier, or filter, of centrosome-nucleated microtubule, blocking the extension of many centrosome-nucleated microtubules, thereby limiting the number of astral and spindle microtubules. This may limit competition for limited cytoplasmic pools of soluble tubulin, which might only be sufficient for a limited number of astral microtubules to extend all the way to cortex. Likewise, an over-abundance of microtubules might interfere with proper spindle formation or function. Moreover, the termination of peri-centrosomal microtubules at the centriculum would likely result in microtubule catastrophe, creating an environment of high tubulin concentration, as observed by Baumgart et al (Baumgart *et al*., 2019); the free tubulin is then captured by the PCM, which is capable of concentrating tubulin *in vitro* (Woodruff *et al*., 2017). Thus, the barrier properties of the centriculum could explain both the accumulation of short peri-centrosomal microtubules and the accumulation of soluble tubulin at the centrosome.

To date, the only centrosome-adjacent membrane that has been examined at nanometer resolution is the *C. elegans* centriculum. Similar membrane structures, however, have been observed in other cell types (Araújo *et al*., 2023; Bergman *et al*., 2015; Bobinnec *et al*., 2003; Diaz *et al*., 2019; Harris, 1975; Karabasheva & Smyth, 2019; Terasaki, 2000; Waterman-Storer *et al*., 1993). In many cases, centrosome-associated membranes are most prominent in the embryo (e.g. *C. elegans*, *Drosophila* and sea urchin). Early embryos typically contain stores of protein and mRNA that are used in the first few embryonic divisions, before the onset of zygotic expression. The centriculum and other centrosome-associated membranes may help limit the number of astral microtubules under conditions where microtubule nucleating capacity exceeds the needs of the embryo. Given the overall conservation in centrosome and ER-associated proteins, it is possible that centrosome structure and function in additional cell types are also affected by an adjacent ER-derived membrane reticulum. If this is the case, then the repertoire of proteins and processes that affect centrosome function is broader than previously appreciated, and defects in these proteins or processes might be a basis for centrosome-related human disease.

## Materials and Methods

### C. elegans strains

The *C. elegans* strains used in this study were derived from the N2 strain (Bristol; (Brenner, 1974) and are listed in Table S1. Strains were maintained at 20°C on OP50 unless treated with RNAi (see below), using standard methods (Brenner, 1974).

### Manipulations of gene and protein expression

#### Feeding RNAi

*E. coli* RNAi feeding clones against *smd-1* F47G4.1, *klp-7* K11D9.1, *zyg-9* F22B5.7 and *tbg-1* F58A4.8 were from a *C. elegans* RNAi feeding library (Open Biosystems, Huntsville, AL). For *klp-7*, *tbg-1*, *zyg-9* or *smd-1* (control (Golden *et al*, 2009)) feeding RNAi treatments, a 5 ml Luria Broth (LB) with 50 µg/ml ampicillin (Sigma Aldrich, Cat # A9518) were inoculated using 1:100 dilution of an overnight saturated culture (at 37°C) of *E. coli* expressing dsRNA of the gene of interest. Once the culture grew to OD_600_ of around 0.5 (∼ 4 h at 37°C), 0.5 M IPTG (Sigma Aldrich, Cat # I6758, 1 mM final concentration) was added to induce the bidirectional transcription of the relevant gene for another 4 h. The culture was centrifuged at 5000 g for 5 min at room temperature, and the pellet was resuspended in 1 ml of fresh LB + ampicillin (50 μg/ml) media. 200 μl of this culture were spread on each RNAi plate (MYOB with 4 mM IPTG and ampicillin 50 µg/ml). For feeding RNAi treatment, 20-40 L4-stage larvae were transferred to RNAi plates at 20°C, and after 48 h (for *klp-7*), or 24 h (for *tbg-1* and *zyg-9*), the RNAi treated worms were dissected on a glass slide, mounted on a 2% agar pad, and imaged as described below. Control RNAi treatments (*smd-1*) were done for the same amount of time as the experimental ones.

#### dsRNA injection RNAi

RNAi against *spd-5* was done using injection of double stranded RNA (dsRNA). A region of *spd-5* from 500-949 bp, was PCR amplified from N2 genomic DNA using oligonucleotides containing T7 or T3 promoter sequence. The amplified region was gel purified and then reamplified with the same PCR primers. The PCR product was purified using the Qiagen MinElute Reaction Cleanup Kit (Cat # 28206). To prepare the RNA for injection, in vitro RNA synthesis was carried out using MEGAscript™ T7 Transcription Kit (Invitrogen Cat # AM1333; for forward strand) and MEGAscript™ T3 Transcription Kit (Invitrogen Cat # AM1338; for the reverse strand) followed by purification using Phenol:CHCl3:Isoamyl Alcohol (invitrogen Cat # 15593031; 25:24:1, v/v) and precipitated using 100% ethanol. The RNA pellet was dissolved in water. To prepare dsRNA, an equal amount (2 µg/µl) of both the ssRNAs were mixed and incubated at 85°C for 3 minutes in an aluminum heat block incubator followed by slow cooling to room temperature for annealing. Injection of dsRNA was done according to Ohta laboratory protocol (Ohta *et al*., 2021). L4s (15-20 worms) were injected with ∼1 μg/μl dsRNA. These worms were maintained at 16°C for 48 h prior to live imaging of early embryos by confocal microscopy.

#### Auxin-mediated degradation

Auxin mediated degradation of ATLN-1 was done as described previously (Maheshwari *et al*., 2023). The *atln-1* gene was tagged with an auxin-inducible degron tag (atln-1::degron) in cells expressing TIR1, an exogenous F-box protein. Worms were transferred to bacteria seeded indole-3 acetic acid (IAA, Alfa Aesar Cat # A10556) plates (MYOB plates with 4 mM IAA) for ∼20-25 minutes, and embryos were imaged immediately thereafter, as per Zhang et al 2015 (Zhang *et al*, 2015).

### Confocal microscopy

#### Imaging

Images were taken using a Nikon confocal Ti2 with Yokagawa CSU-X1 spinning disk and a photometrix Prime 95B camera using a Nikon water/oil 60X 1.2-NA Apo Plan objective. Images were captured using Nikon Elements software version 5.20.00. For imaging, embryos were mounted on 2% agarose pads prepared in standard M9 buffer. Images were taken at z=1 μm intervals unless otherwise mentioned.

#### Image analysis

All images were analyzed using Fiji (Schindelin *et al*., 2012), http://imagej.nih.gov/ij).

### Measurements

#### Measurement of areas and amount

To compare between structures, images were taken by confocal microscopy as described above, and the comparisons between relevant structures were done at their midplane, where these structures is at its largest cross section. Since the centriculum, peri-centrosomal microtubules and centrosome are symmetrically nested within each other, the midplane of any two structures was at the same image plane. To determine the area of a given structure, the boundaries of structure were roughly traced. In the case of the PCM this involved tracing the outer perimeter of SPD-5. For the centriculum and peri-centrosomal microtubule ring, this involved tracing both the inner and outer perimeters of the ring, excluding nuclear membranes and spindle microtubules, respectively, resulting in a toroidal shape (Fig. S1A). For each traced area, the minimum (min) and mean fluorescent values were determined. Empirically, the most consistently faithful representation of the different structures’ areas was by using the formula (mean+min)/2 to set the lower threshold value, and Max (maximum threshold, found in same measurement results FIJI that also included mean and minimum intensity values) to set the upper threshold. These values were entered into the threshold function of FIJI, resulting in consistently accurate representation of the structure’s area (in the case of the microtubule ring, since the difference in intensity between the ring and its center is only about 15%, this method did not result in a ring but rather a solid round area) (Fig. S1A). Of note, while in the present study the min/mean ratios were determined for each structure in each image, these values are highly consistent, and once determined can be applied to a given structure. For example, the min/mean ratio of the traced centriculum region for mCherry::SP12 were 0.79 ± 0.03 (control RNAi; n=16) and 0.76 ± 0.04 (*klp-7* RNAi; n=16); for GFP:: SP12 they were 0.75 ± 0.05 (control RNAi; n=16) and 0.74 ± 0.03 (*klp-7* RNAi; n=20).

For the PCM and peri-centrosomal microtubules, the area and total fluorescence (Raw Intensity) were measured in the thresholded region using the analyze > measure function in FIJI. For the centriculum, the inner perimeter of the thresholded centriculum was selected using wand tool in FIJI to get the void area. For the area defined by the outer perimeter, we traced the thresholded outer edge of the centriculum thereby determining the area bound by the centriculum’s outer perimeter.

#### Microtubule angle measurement

In a zoomed-in screenshot of a 240-slice-thick section of the tomography data containing the area of interest (Maheshwari *et al*., 2023; Redemann *et al*., 2017) originally from Amira, two straight lines were drawn, marking the approximate interior and exterior edges of the centriculum (Fig. 8A; while the edges of the centriculum are curved, at this resolution the edge approaches a straight line that represents the tangent line of the centriculum edge). Microtubules that crossed the interior edge were traced using the straight line tool in FIJI. If a microtubule continued beyond the exterior boundary line of the centriculum, it was categorized as a “continued MT.” Microtubules that stopped before reaching the exterior boundary were categorized as “stopped MT.” To ensure that microtubules defined as “stopped” did not continue outside the range of the slice that was being analyzed, screenshots were taken of 300 slices above and below the same area of the initial screenshot. These images were used to determine whether the traced microtubules continued in another plane. Those that did not continue on either the above or below slice set were considered stopped.

To measure microtubule angles, the angle between the inner centriculum edge line (the tangent line) and the microtubule line was determined. This angle was then subtracted from 90 degrees. The resulting value represents the angle between the microtubule and the perpendicular to the tangent line, also referred to as the normal line (Fig. 8A).

### Focused-Ion beam scanning electron microscope (FIB-SEM) imaging

The FIB-SEM data used in this study was previously published (Rahman *et al*, 2020). 3D segmentation of centriculum structure were done as previously published (Rahman *et al*., 2020) using Amira 6 (version 3D 2022.2, FEI/Thermo Fisher Scientific) default general scheme to segment selected ROI in a semi-autonomous method. Briefly, first, we selected centriculum membranes using a threshold followed by at least three rounds of manual inspection (slice by slice through X, Y, and Z separate planes) to remove incorrect segments and add unsegmented membranes to the final membrane volume (Maheshwari *et al*., 2023; Rahman *et al*., 2020).

To measure the size of centriculum openings (namely open space between reticular membranes that extend all the way through the centriculum), we placed fiducials (on the same plane) next membrane edges on the pronuclear and cortical size of the centriculum. We took TIFF screenshots of the centriculum membrane from behind a central fiducial (i.e., equal distance from the pronuclear and cortical centriculum membrane fiducials) placed on the same plane as perinuclear/cortical fiducials. For each TIFF image, we determined a scale bar separately by marking 11 consecutive pixels as 100 nm. We measured the size of each hole using the threshold-based selection and area measurement tool in Fiji (version 2.14.0/1.54f). To ensure that the centriculum segments in these images are *en face*, we restricted the angle between the center plane and the top and bottom of each image to less than 30°. The actual angle was determined by drawing straight lines from the central fiducial to fiducials that were placed on the top bottom edges of each image. Next, we took TIFF images of the perpendicular view of the above-mentioned lines and utilized the angle measurement tool in Fiji.

### Statistical analysis

All analyses were done using GraphPad Prism (GraphPad Prism [Version 10.1.1 (270)]). For comparisons between two normally distributed samples, unpaired t-tests were used; otherwise, a two-tailed Mann-Whitney test was used as a non-parametric test. For comparison between multiple datasets, one-way ANOVA non-parametric Kruskal-Wallis test with correction for multiple comparisons was used. The statistical tests used, the number of samples analyzed, and the p-values are indicated in the Figure legends. The criterion for statistical significance was set at p < 0.05. Results in all graphs are represented as mean ± standard deviation unless indicated otherwise. The binned frequency distribution graphs in Fig. 7, 8 and Supplemental Figure S3 were done using Microsoft excel. The bin width was set at 500 nm^2^ up to 1000 nm^2^ then from 1001 to 10000 the data were binned at every 1000 nm^2^ with overflow bin set at >10000 nm^2^.

### AI use

AI was not used in any part of this study or manuscript preparation.

## Acknowledgments

We thank Karen Oegema (UC San Diego), Jessica Feldman (Stanford University), Jon Audhya (University of Wisconsin-Madison) and the *Caenohrabditis* Genetics Center for strains. We thank Kevin O’Connell (NIH, NIDDK), Antonina Roll-Mecak (NIH, NINDS), Gunar Fabig (University of Dresden) and Alexey Khodjakov (Wadsworth Center, New York State Department of Health) for helpful discussions. We also thank Kevin O’Connell for comments on the manuscript. We thank Kedar Narayan for support in obtaining FIB-SEM data. R.M., M.M.R., A.R., S.D., R.S.M. and O.C-F. were supported by an NIDDK intramural grant to O.C-F., DK069012-18.

This research was supported by the Intramural Research Program of the National Institute of Diabetes and Digestive and Kidney Diseases (NIDDK) within the National Institutes of Health (NIH). The contributions of the NIH authors were made as part of their official duties as NIH federal employees, are in compliance with agency policy requirements, and are considered Works of the United States Government. However, the findings and conclusions presented in this paper are those of the authors and do not necessarily reflect the views of the NIH or the U.S. Department of Health and Human Services

## Author contribution

The premise of this study was conceptualized by R.M., M.M.R and O.C-F. Experiments were designed, executed and analyzed by R.M., M.M.R, A.R., S.D. R.S.M, and O.C-F. Methodology was developed by R.M., M.M.R, A.R., and S.D. O.C-F supervised the project and wrote the original draft. All authors reviewed and commented on the original draft as well as all other manuscript iterations and accepted the final manuscript.

