## Supplemental Figures for "The centriculum, a membrane reticulum that surrounds *C. elegans* centrosomes, may serve as a microtubule filter"

### Supplemental Figure S1

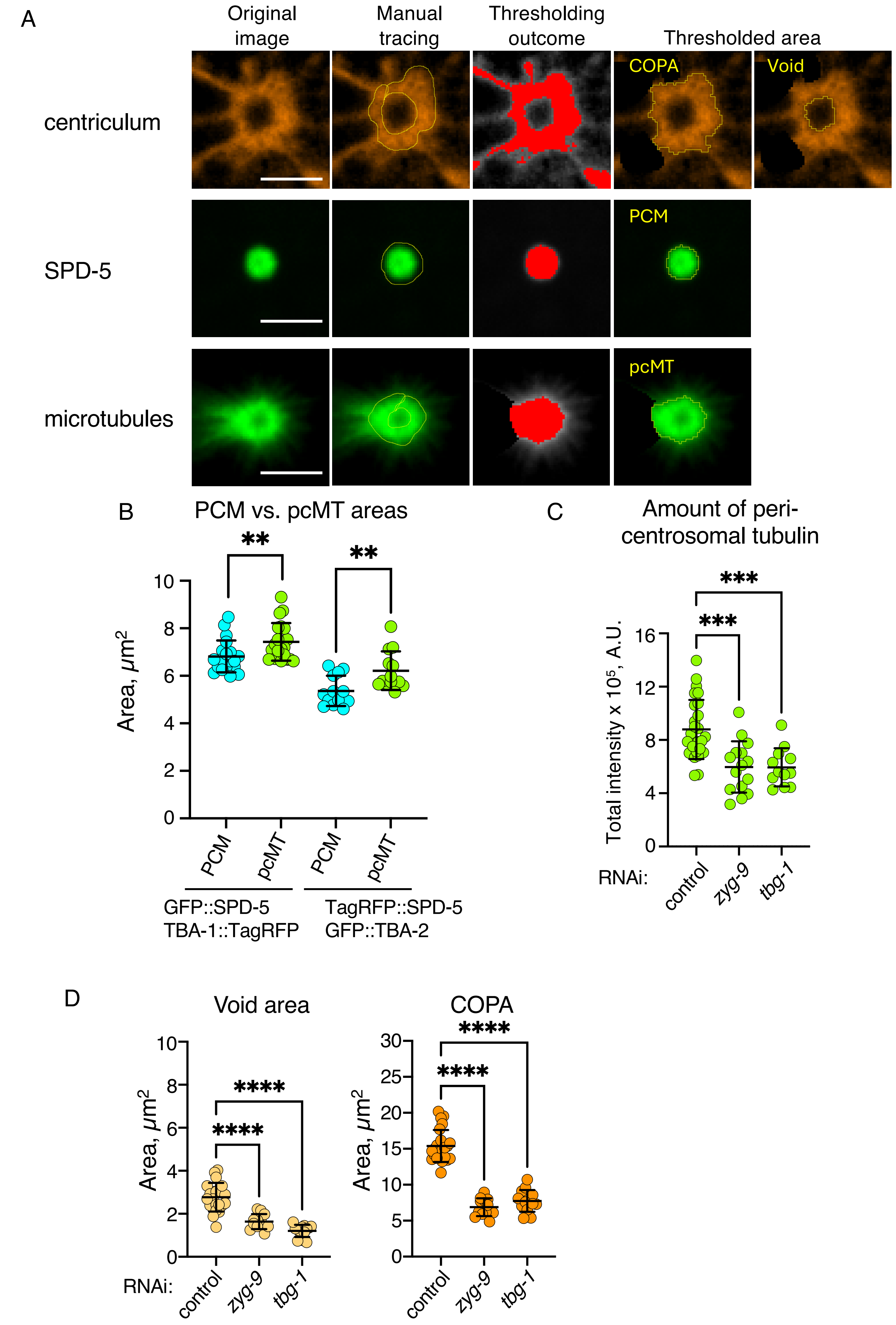

**Supplemental Figure S2**

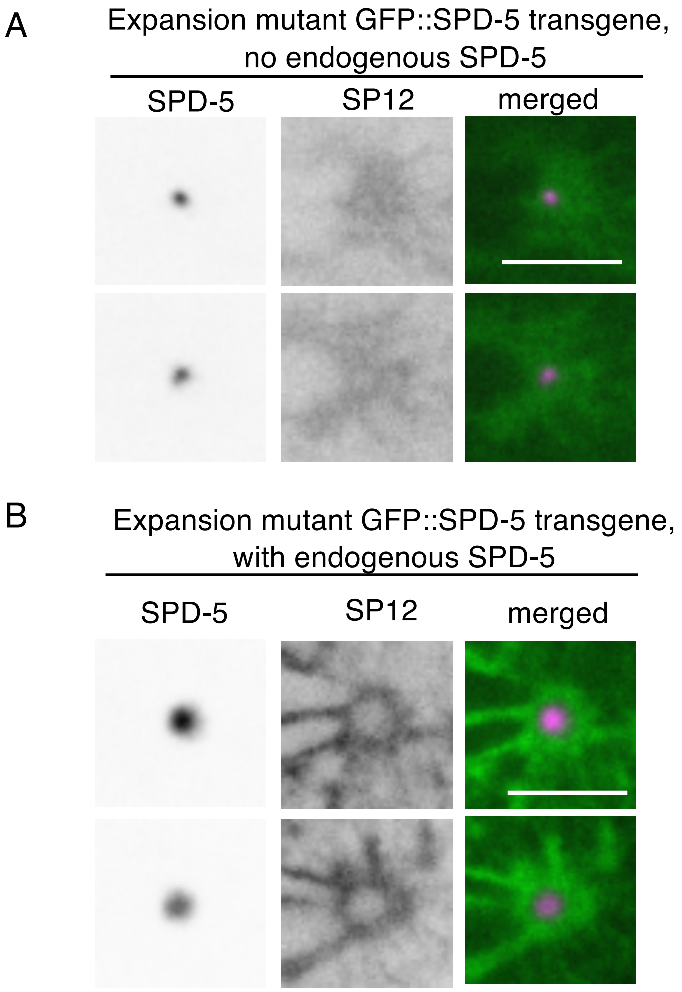

**Supplemental Figure S3**

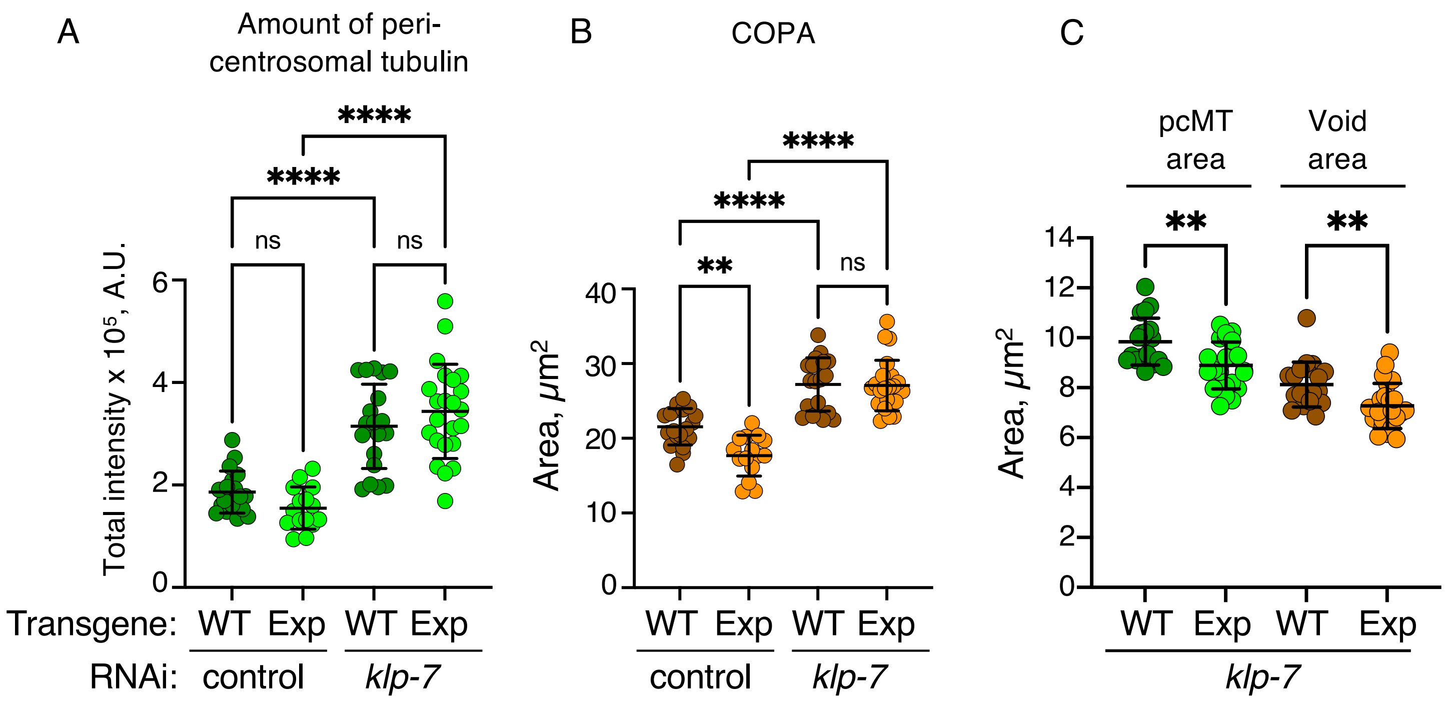

**Supplemental Figure S4**

A. Distribution of GFP-tagged wild type and *spd-5*$^{exp}$ mutant transgenes, in the presence of untagged endogenous *spd-5*:

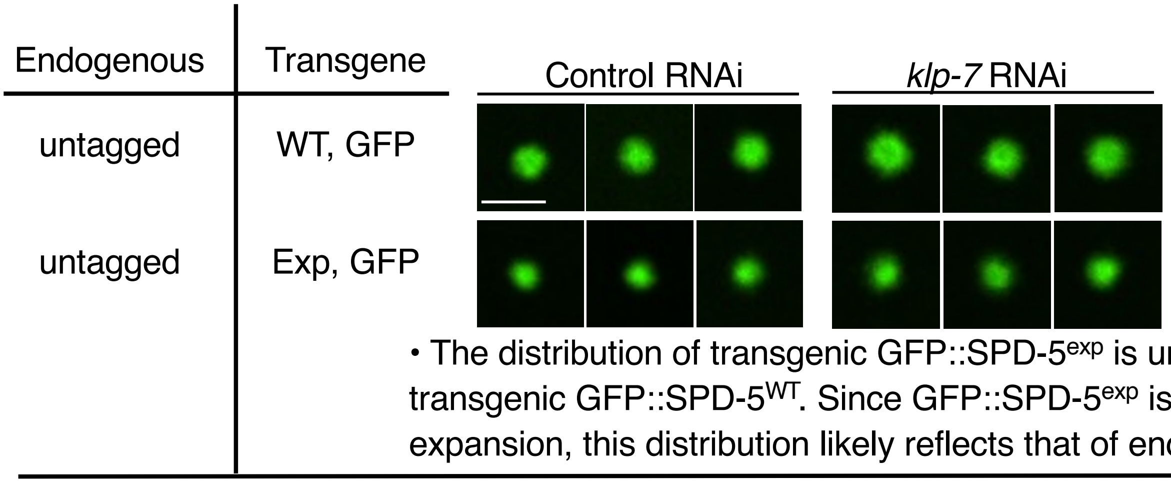


• The distribution of transgenic GFP::SPD-5$^{exp}$ is uniform, similar to transgenic GFP::SPD-5$^{WT}$. Since GFP::SPD-5$^{exp}$ is defective in expansion, this distribution likely reflects that of endogenous SPD-5.

B. Distribution of GFP-tagged wild type and *spd-5*$^{exp}$ mutant transgenes in the presence of GFP-tagged endogenous *spd-5* :

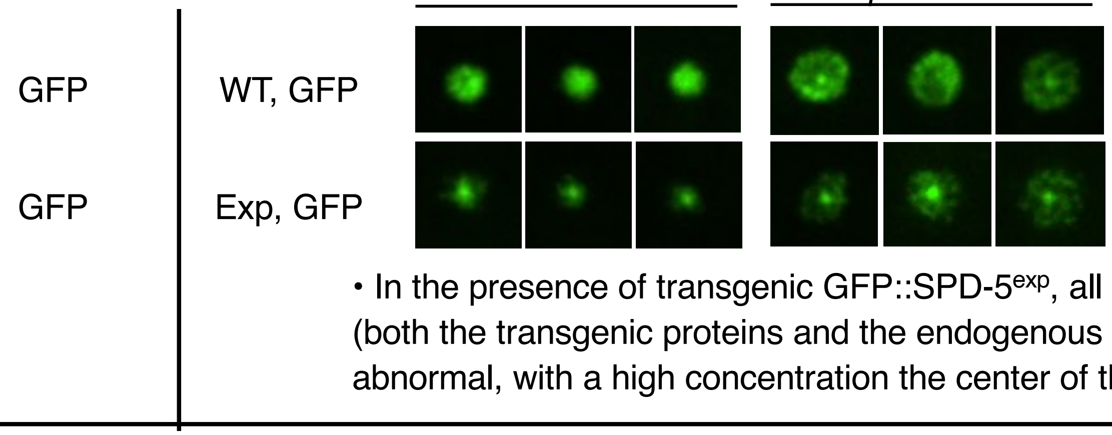


• In the presence of transgenic GFP::SPD-5$^{exp}$, all GFP-tagged proteins (both the transgenic proteins and the endogenous SPD-5) are abnormal, with a high concentration the center of the centrosome.

C. Distribution of GFP-tagged endogenous *spd-5* in the presence of untagged wild type or *spd-5*$^{exp}$ mutant transgenes

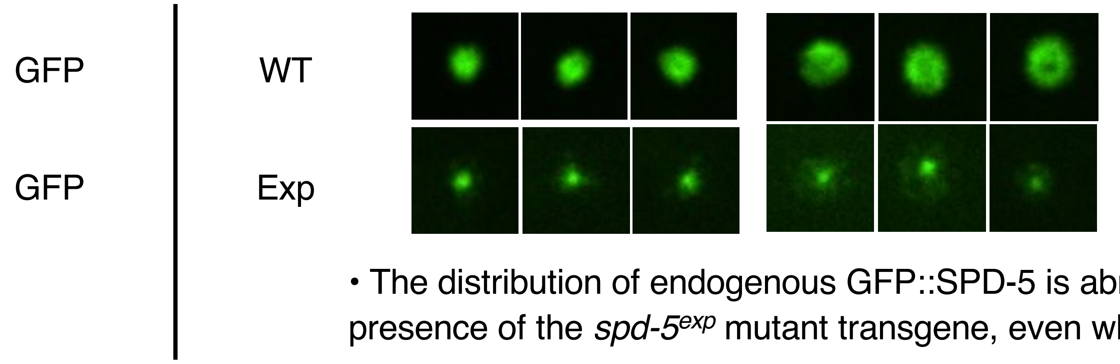


• The distribution of endogenous GFP::SPD-5 is abnormal in the presence of the *spd-5*$^{exp}$ mutant transgene, even when not GFP-tagged.

D. Measurement of endogenously tagged RFP::SPD-5

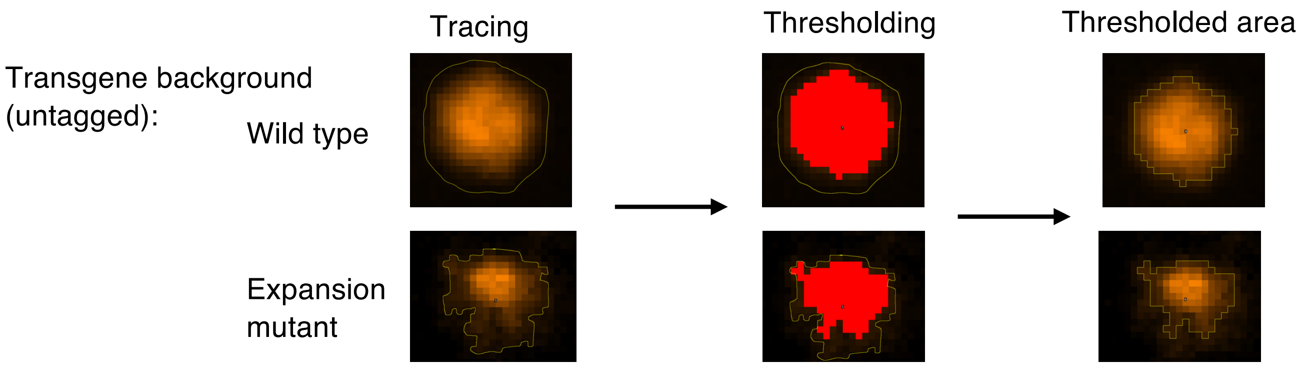

**Supplemental Figure S5**

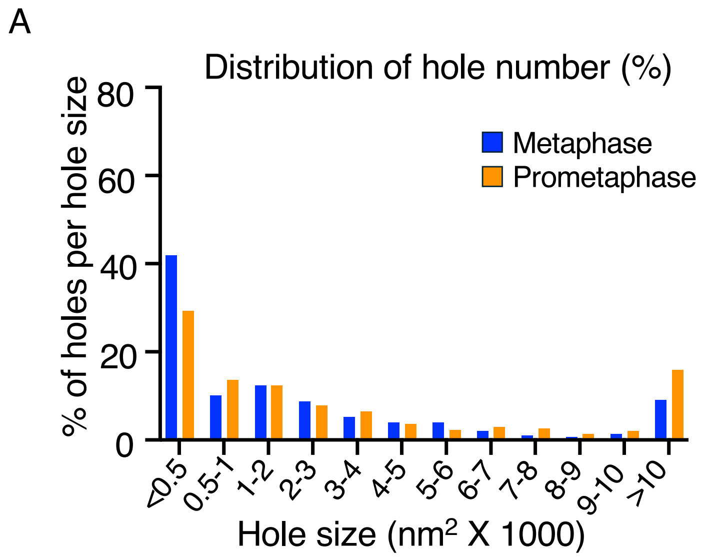
