## Supplemental Material for "The centriculum, a membrane reticulum that surrounds *C. elegans* centrosomes, may serve as a microtubule filter"

**Supplemental Table S1**

| ***C. elegans* strains** | | |
|---|---|---|
| **Name** | **Genotype** | **Source** |
| N2 | wild type (Bristol) | CGC |
| MSN146 | ltIs76 [pAA178: *pie-1p::mCherry::SP12* + *unc-119* (+)]; ltIs25 [pAZ132; *pie-1p::GFP::tba-2 + unc-119 (+)*];*unc-119(ed3)*III | Jon Audhya lab |
| OCF176 | *spd-5(vie26[GFP::spd-5 +loxP]) I; ocfIs2[pie-1p::mCherry::SP12::pie-1 3'UTR +unc119 (+)];his-72(erb77[his-72::linker::mTurquoise2]III; unc-119(ed3)III* | (Maheshwari *et al.*, 2023) |
| OCF181 | *ocfIs2[pie-1p::mCherry::SP12::pie-1 3'UTR +unc119 (+)]; ltIs25 [pAZ132; pie-1p::GFP::tba-2 + unc-119 (+)];his-72(erb77[his-72::linker::mTurquoise2]III; unc-119(ed3)III* | (Maheshwari *et al.*, 2023) |
| OCF183 | *ocfIs2[pie-1p::mCherry::SP12::pie-1 3'UTR +unc119 (+)];ltIs25 [pAZ132; pie-1p::GFP::tba-2 + unc-119 (+)]; his-72(erb77[his-72::linker::mTurquoise2]III; ocf101[atln-1::3xFLAG::degron] IV (CRISPR); TIR1::mRuby IV; unc-119(ed3)III* | (Maheshwari *et al.*, 2023) |
| OCF187 | *ocfIs2[pie-1p::mCherry::SP12::pie-1 3'UTR +unc119 (+)]; cb-unc-119(+)]I; ltSi1141[pOD1021/pVV103; spd-2p::GFP::spd-5 reencoded; cb-unc-119(+)]II; unc-119(ed3)III* | This study |
| OCF189 | *ocfIs2[pie-1p::mCherry::SP12::pie-1 3'UTR +unc119 (+)]; ltSi592[pVV168; spd-2p::GFP-spd-5 mut 653,658, reencoded; cb-unc-119(+)]II; unc-119(ed3)III* | This study |
| OCF193 | *ojIs23 [pie-1p::GFP::SP12 + unc-119(+)]; GFP::PH; tba-1(pg77[tba-1::TagRFP-T + loxP]) I; unc-119(ed3)III* | This study |
| OCF200 | *ltSi592[pVV168; spd-2p::GFP::spd-5 mut 653,658, reencoded; cb-unc-119(+)]II; spd-* | This study |

| | | |
|---|---|---|
| | *5(wow36[tagRFP-T^3xmyc::spd-5])I; unc-119(ed3)III;* | |
| OCF201 | *cb-unc-119(+)]I; ltSi1141[pOD1021/pVV103; spd-2p::GFP::spd-5 reencoded; cb-unc-119(+)]II; spd-5(wow36[tagRFP-T^3xmyc::spd-5])I; unc-119(ed3)III* | This study |
| OCF212 | *tba-1(pg77[tba-1::TagRFP-T + loxP]) I; spd-2p::GFP::spd-5 reencoded; cb-unc-119(+)]II; unc-119(ed3) III* | This study |
| OCF213 | *tba-1(pg77[tba-1::TagRFP-T + loxP]) I; ltSi592[pVV168; spd-2p::GFP::spd-5 mut 653,658, reencoded; cb-unc-119(+)]II; unc-119(ed3)III* | This study |
| OCF214 | *ojIs23 [pie-1p::GFP::SP12 + unc-119(+)];tba-1(pg77[tba-1::TagRFP-T + loxP]) I; ltSi1129[pZZ2; spd-2p::spd-5 (re-encoded);cb-unc-119(+)]II; unc-119(ed3)III* | This study |
| OCF215 | *ojIs23 [pie-1p::GFP::SP12 + unc-119(+)]; tba-1(pg77[tba-1::TagRFP-T + loxP]) I; ltSi1219[pMO104; spd-2p::spd-5 S653A S658A::spd-5 3'UTR; cb-unc119(+)]II; unc-119(ed3)III* | This study |
| OCF218 | *ltSi1129[pZZ2; spd-2p::spd-5 (re-encoded);cb-unc-119(+)]II; spd-5(vie26[GFP::spd-5 +loxP]) I; ocfIs2[pie-1p::mCherry::SP12::pie-1 3'UTR +unc119 (+)]; unc-119(ed3)III* | This study |
| OCF221 | *cb-unc-119(+)]I; ltSi1141[pOD1021/pVV103; spd-2p::GFP::spd-5 reencoded; cb-unc-119(+)]II; spd-5(vie26[GFP::spd-5 +loxP]) I; ocfIs2[pie-1p::mCherry::SP12::pie-1 3'UTR +unc119 (+)];unc-119(ed3)III* | This study |
| OCF223 | *ltSi592[pVV168; spd-2p::GFP::spd-5 mut 653,658, reencoded; cb-unc-119(+)]II; spd-5(vie26[GFP::spd-5 +loxP]) I; ocfIs2[pie-1p::mCherry::SP12::pie-1 3'UTR +unc119 (+)];unc-119(ed3)III* | This study |
| OCF227 | *ltSi1219[pMO104; spd-2p::spd-5 S653A S658A::spd-5 3'UTR;cb-unc-119(+)]II; spd-5(vie26[GFP::spd-5 +loxP]) I; ocfIs2[pie-* | This study |

| | | |
|---|---|---|
| | *1p::mCherry::SP12::pie-1 3' UTR +unc119 (+)],unc-119(ed3)III* | |
| OCF233 | *ojIs23 [pie-1p::GFP::SP12 + unc-119(+)]; ltSi1219[pMO104; spd-2p::spd-5 S653A S658A::spd-5 3'UTR;cb-unc-119(+)]II; spd-5(wow36[tagRFP-T^3xmyc::spd-5])I; unc-119(ed3)III* | This study |
| OCF234 | *ojIs23 [pie-1p::GFP::SP12 + unc-119(+)]; ltSi1129[pZZ2; spd-2p::spd-5 (re-encoded);cb-unc-119(+)]II; spd-5(wow36[tagRFP-T^3xmyc::spd-5])I; unc-119(ed3)III* | This study |
| OCF240 | *spd-5(wow36[tagRFP-t^3xmyc::spd-5]) I; ltIs25 [pAZ132; pie-1p::GFP::tba-2 + unc-119 (+)]* | This study |
| OCF247 | *spd-5(wow52[GFP^3xflag::spd-5]) I tba-1(pg77[tba-1::TagRFP-T + loxP]) I* | This study |

### Supplemental Figure legends

#### Supplemental Figure S1:

**A)** Images of a centriculum, PCM and microtubules of a 1-cell stage metaphase embryos demonstrating how manual tracing and thresholding is used to measure these structures' areas. See the Material and Methods section for the detailed process. Scale bar=5 μm.

**B)** Comparison of PCM and pcMT areas using different fluorescent proteins pairs: OCF247: GFP::SPD-5; TBA-1::tagRFP (same data as shown in Fig 1E, n=20, p=0.0043); and OCF240: RFP::SPD-5; GFP::TBA-2) n=13, p=0.0089, using the Mann-Whitney test.

**C)** Quantification of the amount of peri-centrosomal tubulin fluorescence for the images such as shown in Fig 2A, using strain OCF181. n= 26, 15, and 12 for control, *zyg-9* and *tbg-1* RNAi treatments, respectively. p=0.0001 for control vs. *zyg-9* and p=0.0003 for control vs. *tbg-1* RNAi treatments using ordinary one-way ANOVA. Error bars represent mean and standard deviation.

**D)** Quantification of the void area and COPA for the images such as shown in Fig 2D, using strain OCF176. For void area: n= 22,12, and 14 for control, *zyg-9* and *tbg-1* RNAi treatments, respectively. p<0.0001 for control vs. *zyg-9* and for control vs. *tbg-1* RNAi treatments using ordinary one-way ANOVA. For COPA: n= 23,12, and 14 for control, *zyg-9* and *tbg-1* RNAi treatments, respectively. p<0.0001 for control vs *zyg-9* and for control vs *tbg-1* RNAi treatments using ordinary one-way ANOVA. Error bars represent mean and standard deviation. Note that the data for the control RNAi are the same as shown in Fig 1D.

#### Supplemental Figure S2:

**A and B)** Additional examples of centricula and PCM from 1-cell embryos at metaphase expressing mCherry::SP12 (green in merged images) and transgenic GFP::SPD-5$^{exp}$ (magenta in merged images) without (panel A) or with (panel B) endogenously expressed SPD-5 (OCF 189), as also shown in Fig 3C and D. Endogenous *spd-5* was down-regulated in panel A using RNAi. Scale bar=5 μm.

**Supplemental Figure S3:**

**A and B)** Quantification of peri-centrosomal tubulin amount and COPA from the control and *klp-7* RNAi treated embryos as shown in Fig 5B (strains OCF214 and 215). n=19 and 15 for wild type and *spd-5*$^{exp}$ transgenes, respectively, treated with control RNAi. n=20 and 23 wild type and *spd-5*$^{exp}$ transgenes, respectively, treated with *klp-7* RNAi. p-values, determined by ordinary one-way ANOVA, are (from left to right): panel A: 0.5932, <0.0001, <0.0001 and 0.5529; panel B: 0.0012, <0.0001, <0.0001 and 0.8739. Error bars represent mean and standard deviation.

**C)** Quantification of pcMT area and void area following *klp-7* RNAi treatment as shown in Fig 5B. n=20 and 23 for wild type and *spd-5*$^{exp}$ transgenes, respectively. p-values are 0.0021 (pcMT area) and 0.0061(void area) as determined by ordinary one-way ANOVA. Error bars represent mean and standard deviation.

**Supplemental Figure S4:**

**A-C)** A comparison of endogenous and/or transgenic *spd-5* distribution when fused to GFP, in the presence of untagged or wild type or *spd-5*$^{exp}$ transgenes. Images are of three different centrosomes following control or *klp-7* RNAi treatment. The latter allows for more detailed protein distribution. The strains used were as follows: panel A: OCF187 and OCF189, expressing the wild type or *spd-5*$^{exp}$ transgene, respectively, both tagged with GFP and expressing untagged endogenous *spd-5*; panel B: OCF221 and OCF223, expressing the wild type or *spd-5*$^{exp}$ transgene, respectively, both tagged with GFP and expressing GFP-tagged endogenous *spd-5*; and panel C: OCF218 and OCF227, expressing the untagged wild type or *spd-5*$^{exp}$ transgene, respectively, and GFP-tagged endogenous *spd-5*. Scale bar=5 μm.

**D)** Steps to quantify PCM area of centricula of 1-cell embryos expressing RFP::SPD-5 with either transgenic wild type *spd-5* (OCF234) or untagged *spd-5*$^{exp}$ mutant (OCF233). See Materials and Methods for detailed quantification process.

**Figure S5:**

**A)** Binned frequency distribution of holes present on the pronuclear side of metaphase (in blue) and prometaphase (orange) centricula from 1-cell embryos. The metaphase data are the same as shown in Fig 7D. Bin size range is shown on the x-axis. n=308 for prometaphase holes; pooled from 6 images taken from 3 centricula (2 images per centriculum).

**B)** Binned frequency distribution of the percentage of total open area per hole size range, for holes in metaphase (in blue) and prometaphase (orange) centricula from 1-cell embryos, using the same data as in panel A. The data for metaphase centricula are the same as shown in Fig 7E.
